# High-throughput mapping of single neuron projections by sequencing of barcoded RNA

**DOI:** 10.1101/054312

**Authors:** Justus M Kebschull, Pedro Garcia da Silva, Ashlan P Reid, Ian D Peikon, Dinu F Albeanu, Anthony M Zador

## Abstract

Neurons transmit information to distant brain regions via long-range axonal projections. In the mouse, area-to-area connections have only been systematically mapped using bulk labeling techniques, which obscure the diverse projections of intermingled single neurons. Here we describe MAPseq (Multiplexed Analysis of Projections by Sequencing), a technique that can map the projections of thousands or even millions of single neurons by labeling large sets of neurons with random RNA sequences ("barcodes"). Axons are filled with barcode mRNA, each putative projection area is dissected, and the barcode mRNA is extracted and sequenced. Applying MAPseq to the locus coeruleus (LC), we find that individual LC neurons have preferred cortical targets. By recasting neuroanatomy, which is traditionally viewed as a problem of microscopy, as a problem of sequencing, MAPseq harnesses advances in sequencing technology to permit high-throughput interrogation of brain circuits.

## Introduction

Neurons transmit information to distant brain regions via long-range axonal projections. In some cases, functionally distinct populations of neurons are intermingled within a small region. For example, nearby hypothalamic nuclei regulate basic drives including appetite, aggression, and sexual attraction (Kennedy et al., 2014; Sternson, 2013), and neurons from these nuclei project to distinct targets. In visual cortical area V1, responses to visual stimuli are matched to the properties of the higher visual areas to which the neurons project (Glickfeld et al., 2013; Movshon and Newsome, 1996). Findings such as these suggest that the information transmitted by individual neurons may be tailored to their targets. Such selective routing of information requires an anatomical substrate, but there is currently no high-throughput method for determining the diverse projection patterns of individual neurons.

At present, there is a steep tradeoff between throughput and resolution in anatomical approaches to mapping long-range connections. In conventional anterograde brain mapping studies, a fluorescent or enzymatic label is used to enable visualization of cell bodies and distal projections by lightmicroscopy. Bulk techniques query the projections of many neurons labeled at a single injection site and thus sample the aggregate architecture of an entire neuronal population. There have been several large-scale efforts,including the Allen Brain Projection Atlas (Oh et al., 2014) and the iConnectome (Zingg et al., 2014), to systematically map mesoscopic connectivity. Although fast, such bulk methods obscure the diversity of the many projection neurons labeled in any one experiment. Consider, for example, a single source area that projects to three downstream areas (Fig 1a). This projection pattern enables neurons in the source area to send information to the three downstream areas. However, identical bulk projection patterns could arise in multiple ways: (i) from a one-to-one architecture, in which each neuron targets only a single downstream area (*left*); (ii) from an all-to-all architecture, in which each neuron targets every downstream area (*middle*); or (iii) from a host of more complicated architectures (*right*). With conventional bulk labeling these three projection patterns, which have different functional implications, are indistinguishable without further experimentation.

Several alternative methods have been developed to complement conventional anterograde bulk-labeling approaches. For example, genetically defined subpopulations of neurons within an area can be targeted by expressing a marker such as Cre recombinase (Gong et al., 2007; Harris et al., 2014; Huang, 2014). Similarly, subpopulations can be defined using retrograde (Lima et al., 2009; Oyibo et al., 2014; Wickersham et al., 2007a) or trans-synaptic viruses (DeFalco et al., 2001; Wickersham et al., 2007b). However, because such approaches rely on positing defined subpopulations, they cannot easily be used to screen for the myriad possible complex projection patterns neurons might exhibit.

The most general and unbiased approach to distinguishing among the architectures in Fig. 1a relies on single neuron anterograde tracing. Current methods for achieving single neuron resolution require individually labeling one or, at most, a few cells per brain (Economo et al., 2016), a labor-intensive approach that affords highresolution at the cost of low-throughput. Although single-neuron tracing can be multiplexed by labeling individual neurons with different colored fluorophores (Ghosh et al., 2011; Livet et al., 2007), in practice the extent of multiplexing is limited by the number of colors — at most 5-10 — that can be resolved by microscopy.

**Figure. 1:**
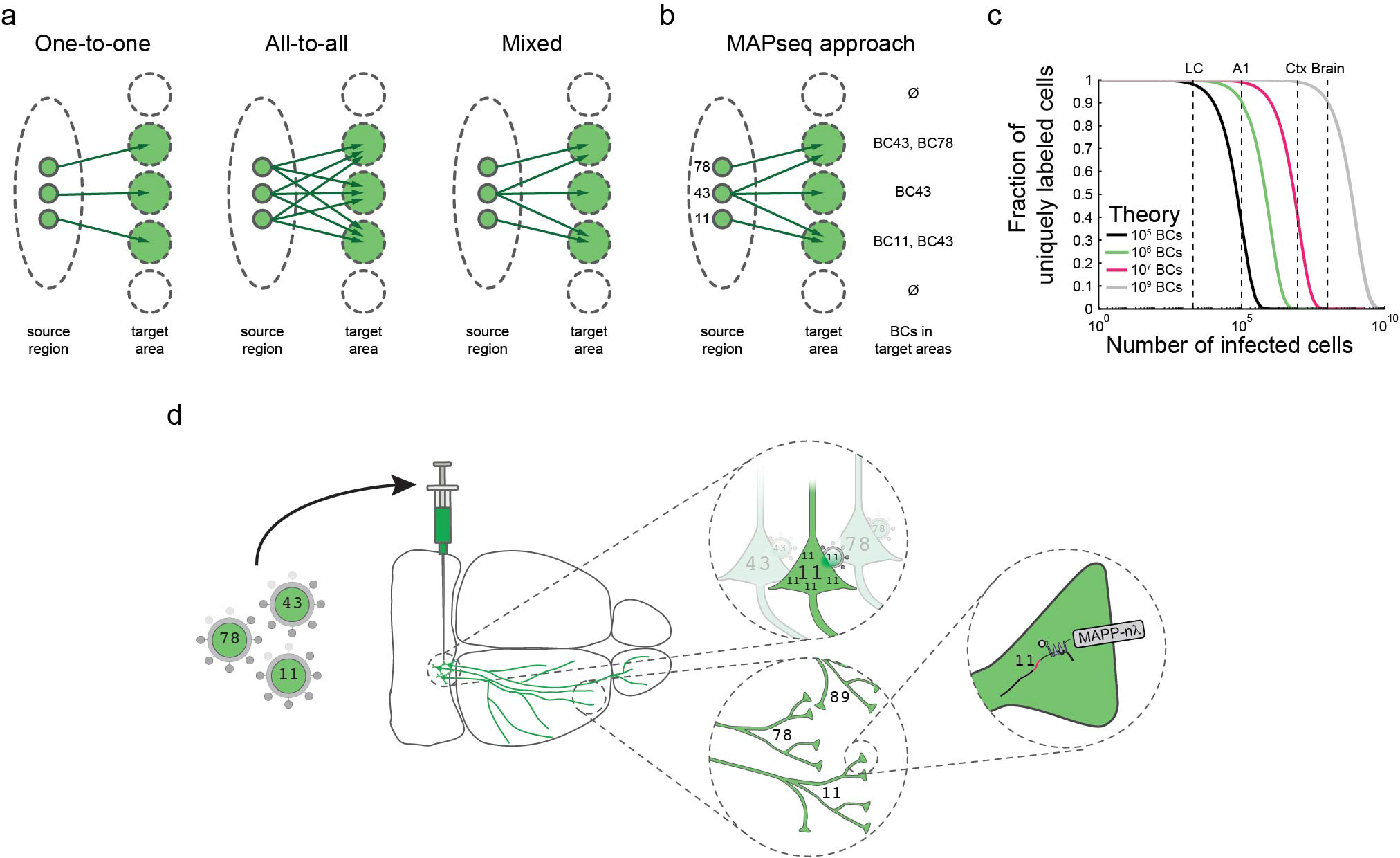
Barcoding allows high-throughput single neuron tracing. (a) Identical bulk mapping results can arise from different underlying projection patterns. (b) Single neuron resolution can be achieved by randomly labeling neurons with barcodes and reading out barcodes in target areas. (c) The expected fraction of uniquely labeled cells is given by *F*=(1-1/*N*)^(*k*-1)^, where *N* is the number of barcodes and *k* is the number of infected cells, assuming a uniform distribution of barcodes. The number of neurons for various mouse brain areas are indicated according to refs (Herculano-Houzel et al., 2006; Schüz and Palm, 1989) (A1= primary auditory cortex; Ctx = neocortex). (d) In MAPseq, neurons are infected at low multiplicity of infection (MOI) with a barcoded virus library. Barcode mRNA is expressed, trafficked and can be extracted from distal sites as a measure of single neuron projections.

Here we describe MAPseq, a novel approach in which the speed and parallelization of high-throughput sequencing is exploited for brain mapping. MAPseq achieves multiplexing by using short, random RNA barcodes to uniquely label individual neurons (Mayer et al., 2015; Walsh and Cepko, 1992; Zador et al., 2012) (Fig 1b). The key advantage of using barcodes is that their diversity grows exponentially with the length of the sequence, overcoming the limited diversity of the resolvable color space. The pool of unique barcode identifiers is effectively infinite; even a 30 nucleotide (nt)-sequence has a potential diversity of 4^3^≈10^18^ barcodes, far surpassing the ~10^8^ neurons in the mouse brain (Herculano-Houzel et al., 2006). Because high-throughput sequencing can quickly and inexpensively distinguish these barcodes, with MAPseq we can potentially read out the projections of thousands or even millions of individual neurons in parallel within a single brain (Fig. 1c).

In MAPseq, we uniquely label neurons in a source region by injecting a viral library encoding a diverse collection of barcode sequences. The barcode mRNA is expressed at high levels and transported into the axon terminals at distal target projection regions (Fig. 1d). To read out single neuron projection patterns, we then extract and sequence barcode mRNA from the injection site, as well as from each target region of interest. Spatial resolution of MAPseq is limited mainly by the precision of target dissection. Although MAPseq, like GFP tracing, does not distinguish fibers of passage, we minimize their contribution by avoiding large fiber bundles during the dissection of target areas. Using this procedure, the brain-wide map of projections from a given area can be determined in less than a week. By reformulating projection mapping as a problem of sequencing, MAPseq harnesses advances in high-throughput sequencing to permit efficient single neuron circuit tracing.

## Results

As a proof-of-principle, we applied MAPseq to LC, a small nucleus in the brainstem that is the sole source of noradrenaline to the neocortex (Foote and Morrison, 1987). Early bulk tracing experiments revealed that the LC projects broadly throughout the ipsilateral hemisphere, leading to the view that the LC broadcasts a generalized signal that modulates overall behavioral state (Foote and Morrison, 1987; Foote et al., 1983; Loughlin et al., 1982; Waterhouse et al., 1983). This view has recently been supported by more sophisticated retrograde bulk tracing experiments, which reinforce the idea that LC neurons project largely indiscriminately throughout the entire ipsilateral hemisphere (Schwarz et al., 2015). However, other reports have challenged this model. Using double retrograde labeling methods, these experiments uncovered separate populations of LC neurons projecting to different areas of cortex (Chandler et al., 2014; Chandler and Waterhouse, 2012), raising the possibility that the LC exerts differential control over different cortical areas. To address this controversy, we applied MAPseq to LC to obtain a large number of projection patterns at single neuron resolution.

In what follows we first show that long-range projections of neurons can be efficiently and reliably determined using barcode mRNAs, the abundance of which we interpret, like GFP intensity, as a quantitative measure of projection strength. Next, we establish the theoretical and practical foundations of randomly labeling large numbers of neurons with a viral barcode library, critical for ensuring single cell resolution for MAPseq. We then apply MAPseq to the LC, and find that individual neurons have a variety of idiosyncratic projection patterns. Some neurons project almost exclusively to a single preferred target in the cortex or olfactory bulb, whereas others project more broadly. Our findings are consistent with, and reconcile, previous seemingly contradictory results about LC projections. Finally we show that MAPseq can be multiplexed to two and potentially many injections in the same animal, which will allow the projection patterns from many brain areas to be determined efficiently and in the same brain without the need for registration across animals.

### Using RNA to trace neurons

Conventional neuroanatomical tracing methods rely on filling neurons with dyes or proteins, so that neural processes can be resolved by microscopy. An implicit assumption of these techniques—albeit one that has rarely been rigorously tested—is that the tracer fills the neuron abundantly and uniformly, so that the strength of the signal corresponds to the quantity of labeled neural process, independent of distance from the soma. For barcode mRNAs to act as a comparable label in MAPseq, we sought to maximize the abundance and uniformity of barcode mRNA in distal processes. We used two strategies to achieve this goal.

First, we expressed a modified presynaptic protein that was designed to specifically bind to and transport barcode mRNA into axon terminals. We engineered this protein, which we denote MAPP-nλ, as part of a larger project aiming to read out synaptic connectivity using mRNA. To generate MAPP-nλ, we began with pre-mGRASP, a protein engineered to localize at the presynaptic terminal due to fusion with trafficking signals from the endogenous presynaptic protein NRXN1β (Kim et al., 2011). We then inserted four copies of the nλ RNA binding domain (Daigle and Ellenberg, 2007) into the cytoplasmic domain of the protein. Derived from the λ phage λ_N_ protein, the nλ domain is a 22 amino acid peptide which strongly and specifically binds to a 15-nt RNA hairpin, termed boxB. We added four copies of the boxB hairpin to the barcode mRNA, ensuring coupling of MAPP-nλ to the barcode mRNA and thus transport of barcode mRNA into axon terminals (Daigle and Ellenberg, 2007). Second, we delivered the barcode sequence using recombinant Sindbis virus, a virus which can rapidly achieve very high expression levels (Ehrengruber, 2002). We used a novel Sindbis packaging system which, unlike previous systems, is both neurotropic and propagation incompetent (Kebschull et al., 2015) (Supplementary Fig 1a-g; Supplementary Note 1). All components necessary for MAPseq are expressed from a dual promoter virus that generates two subgenomic RNAs (Fig. 2a). The first encodes the MAPP-nλ protein. The second RNA encodes a random 30-nt barcode, as well as the boxB sequence, downstream of a GFP marker (Supplementary Fig. 1h,i). We reasoned that combining these two strategies would maximize our ability to reliably detect barcode mRNA in distal processes.

**Figure 2:**
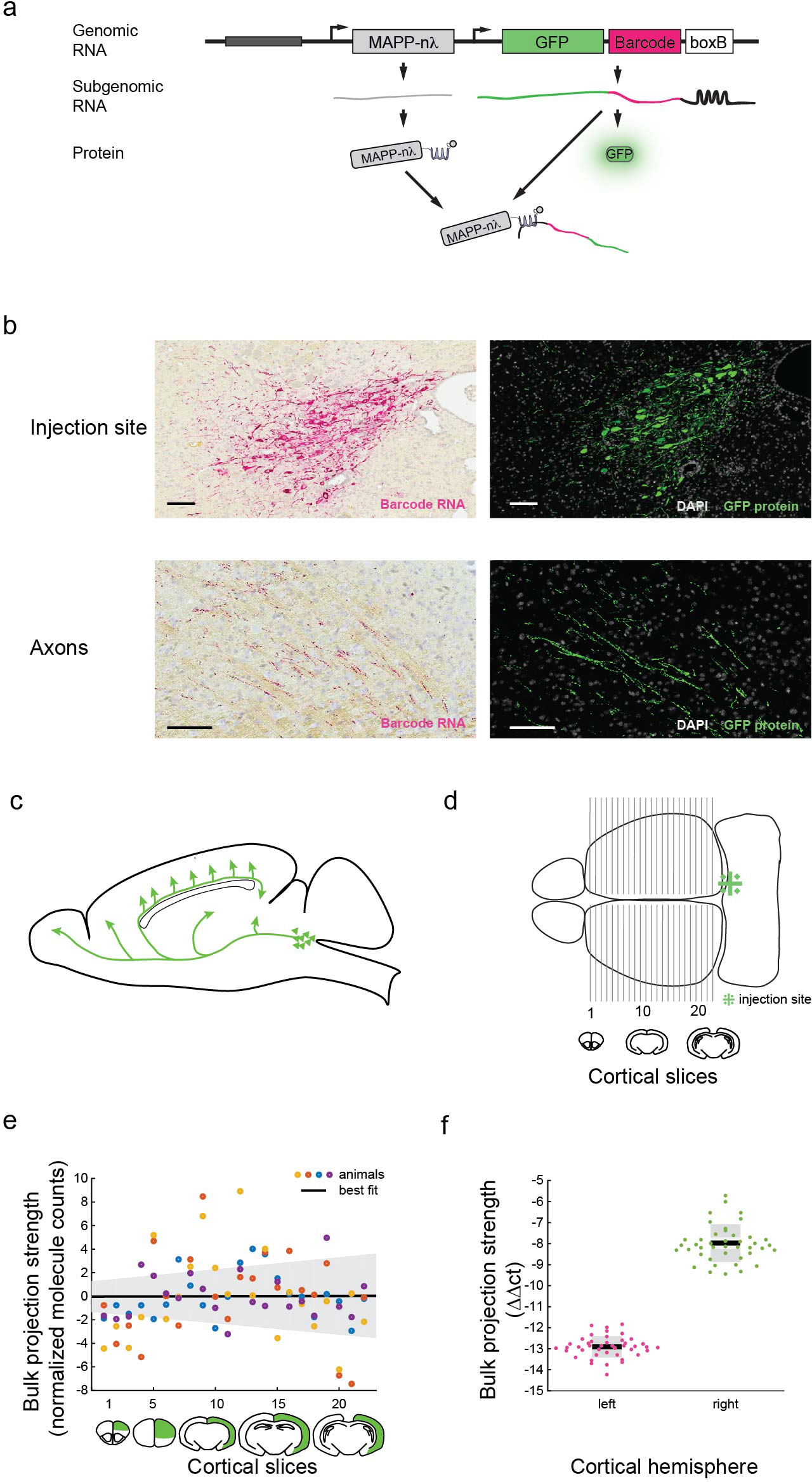
Barcoded Sindbis virus can be used for projection mapping, (a)A dual promoter Sindbis virus was used to deliver barcodes to neurons. The virus encoded GFP, barcodes and MAPP-nλ. (b) Barcode mRNA labeling of LC neurons is comparable to GFP labeling of these neurons in an adjacent 6μm slice both at the injection site (*top*) and in the axon tract (*bottom*). Scale bar =lOOμm. Representative data from 3 animals, (c) Axons from LC project rostrally from the cell body, before changing direction and innervating cortex. LC axons that project to frontal cortices have thus traveled only about half as long as axons innervating visual cortex. (d) We injected right LC with MAPseq virus and dissected cortex along the anterior-posterior axis as shown. (e) Bulk projection strength of LC to ipsilateral cortex as measured by barcode mRNA is independent of the anterior-posterior position of the cortical slice, suggesting a uniform RNA fill of LC axons. N=4. (f) qPCR for barcode mRNA shows approximately 30× stronger LC projections to ipsi-than to contralateral cortex. N=2 animals and 21 cortical slices per animal. BC=barcodes. The y-axis displays ΔΔct values, which are equivalent to the log_2_(foldchange of barcode mRNA per sample) normalized to β-actin levels in each sample andto the amount of barcode mRNA in the injection site of each animal (Livak and Schmittgen, 2001).

We injected barcoded virus into right LC (Supplementary Fig. 1j,k,l) and examined barcode localization by *in situ* RNA hybridization 44 hours after injection. We observed robust barcode mRNA localization in the soma and neuronal processes, in a pattern similar to that of coexpressed GFP (Fig. 2b). This suggested that barcode mRNA could effectively fill local neuronal processes.

To determine whether the barcode mRNA fills distal neuronal processes uniformly, we exploited the particular anatomy of LC projection neurons. LC neurons that project to cortex send their processes all the way to the rostral end of the brain, before changing direction and moving caudally to innervate cortical areas (Fig. 2c). Axons that project to visual cortex are therefore approximately twice as long as those that project to frontal cortices. From bulk tracing studies it is known that LC innervation is homogeneous along the rostro-caudal axis (Schwarz et al., 2015; Waterhouse et al., 1983). Thus if barcode mRNA were not efficiently transported to distal processes, we would expect to find more barcode mRNA in rostral regions of cortex. To assess this, we injected barcoded virus into LC, cut 300μm coronal slices of the entire cortex (Fig. 2d), and quantified the amount of barcode mRNA from each ipsilateral and contralateral slice. Consistent with previous results using GFP and other tracing methods (Schwarz et al., 2015; Waterhouse et al., 1983) we found approximately uniform projections throughout the ipsilateral cortex (p=0.972 F-statistic vs. constant model; Fig. 2e); in particular, we found no evidence that distal processes were more weakly labeled than proximal processes. Also consistent with previous results, we observed much weaker (30.6-fold less; p=4×10^-31^ paired student’s t-test) projections to the contralateral cortex (Fig. 2f). These results suggest that barcode mRNA fills distal and proximal processes with about equal efficacy, so that the barcode mRNA can be interpreted in the same way as the fluorophores and dyes used in conventional tracing studies.

### Unique labeling of neurons with barcodes

In conventional single-neuron tracing, the main challenge to multiplexing is the low diversity of labels (fluorophores or enzymes) available to disambiguate individual neurons. To overcome this challenge, MAPseq labels neurons with short, random RNA barcodes delivered by infection with a diverse viral library. Ideally, each labeled neuron should have exactly one unique barcode. Here we consider the factors that could lead to deviations from this ideal scenario: (i) more than one barcode per neuron (multiple labeling); and (ii) more than one neuron per barcode (nonunique or degenerate labeling). As discussed below, deviations resulting from multiple labeling are much less of a concern than those resulting from degenerate labeling.

A neuron may express more than one barcode if it is infected by more than one viral particle. Such multiple labeling will lead to an overestimate of the number of neurons identified, but will not distort the projection patterns recorded for individual neurons (Fig. 3a). Furthermore, even estimates of the relative abundances of different neuronal classes will, on average, be accurate. Assume for example that two neurons A and B are each labeled with 10 different barcodes. In this scenario, MAPseq will discover 10 instances of neuron A and 10 of B; but even though the absolute number of neurons is incorrect, the facts that neurons A and B have distinct projection patterns, and that these patterns occur in a 1:1 ratio, are accurately inferred. Thus multiple labeling will not, on average, lead to mischaracterization of neuronal classes or of their relative frequency in the population.

**Figure 3:**
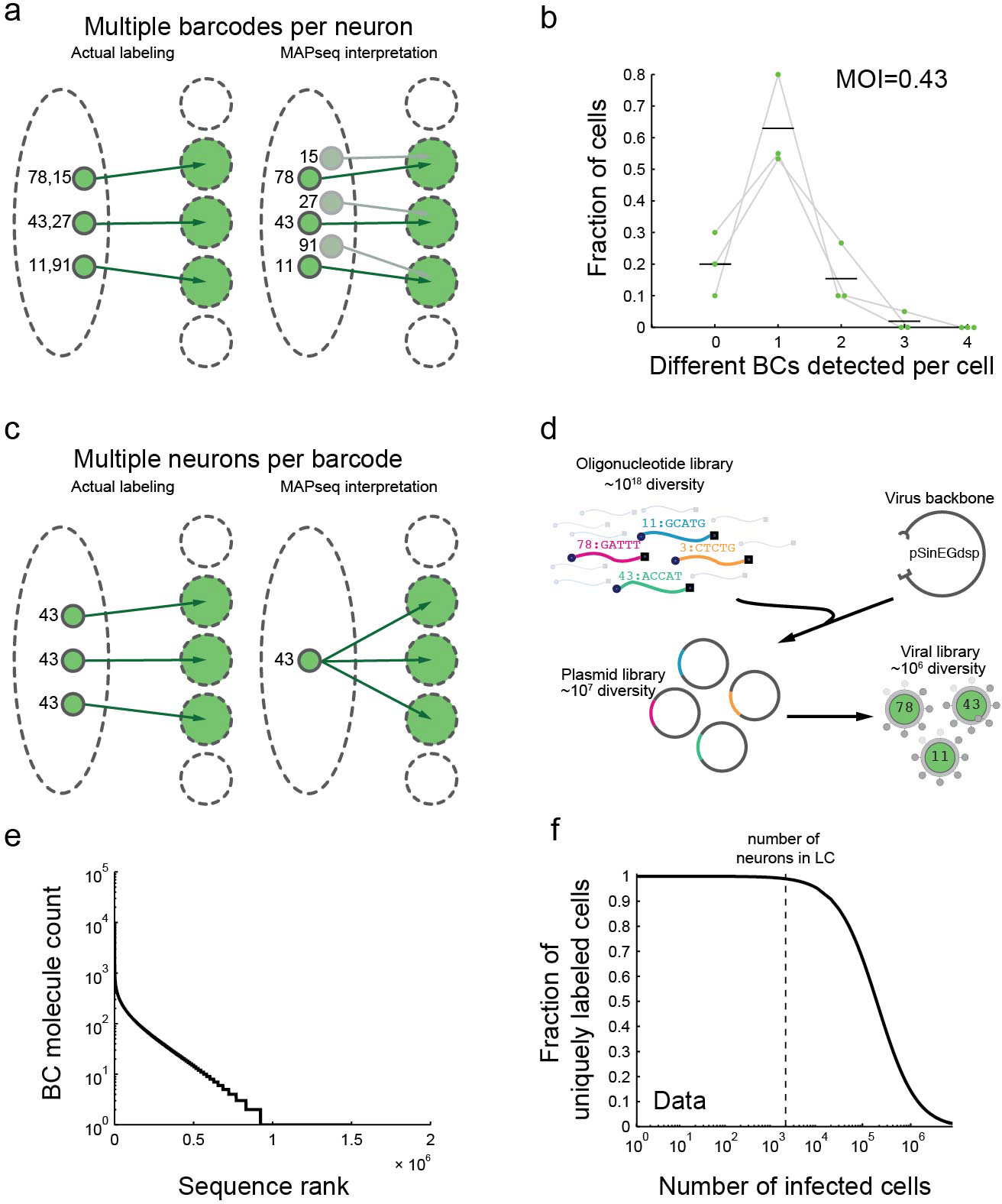
Random labeling of neurons with a barcoded virus library can achieve unique labeling of many neurons, (a) When single neurons are labeled with several barcodes, MAPseq will overestimate of the number of neurons identified, but will not distort the projection patterns recorded for individual neurons, (b) Single cell isolation of GFP-positive, barcoded neurons, followed by sequencing of their barcode complement reveals a low chance of double infection. We interpret neurons for which no barcodes were recovered as technical failures of cell isolation, rather than biological phenomena. N=3 animals. Mean and individual data points are plotted, (c) When several neurons share the same barcode, MAPseq misinterprets this as a single neuron whose projection pattern is given by the union of the projection patterns of the two infected neurons, (d) High diversity Sindbis virus libraries are produced by shotgun cloning random oligonucleotides into a plasmid followed by virus production, (e) The virus library used in this work has a diversity of ~10^6^ different barcodes (BC), but the distribution was non-uniform. The sequence rank is a number that ranges from 1 to the total number of barcodes, where 1 corresponds to the most abundant sequence, 2 to the second most abundant and so on. (f) Based on the empirically observed non-uniform barcode distribution, we determined that the virus library used is sufficiently diverse to uniquely label all of LC with low error rate.

Nevertheless, to simplify the interpretation of MAPseq results we sought to minimize the multiplicity of infection (MOI) by titrating the concentration and volume of virus injected. To estimate the MOI, we isolated a total of 45 individual neurons from 3 animals injected with MAPseq virus into the right LC and sequenced the barcodes within each neuron. On average, infected LC neurons contained 1.2+/-0.1 barcodes each, implying MOIof 0.43 (Fig. 3b). Only 21+/-11% of neurons contained more than one barcode, with most of these neurons carrying two barcode sequences and only 1.7+/-2.9% of neurons containing three barcode sequences.

The second deviation from the ideal scenario is non-unique labeling. If two neurons share the same barcode, then MAPseq misinterprets this as a single neuron whose projection pattern is given by the union of the projection patterns of the two infected neurons (Fig. 3c). The probability that two neurons are infected by the same barcode depends on the number of infected cells relative to the number of available barcodes. Trivially, if the number of infected cells is larger than the number of available barcodes, unique labeling of all neurons cannot be achieved. Conversely, if the number of available barcodes is much higher than the number of infected cells, every neuron will be labeled with a different barcode purely by chance.

To determine whether our barcode diversity was sufficient to ensure unique labeling, we first quantified the number of neurons. We counted 1985+/-132 (N=6 animals) neurons expressing tyrosine hydroxylase, a noradrenergic marker. The size of this neuronal population is approximately 15 orders of magnitude smaller than the theoretical diversity (4^3^≈10^18^) of a library of 30-nucleotide barcodes, so in theory unique labeling would be virtually certain. In practice, however, the actual diversity of a virus library is limited by bottlenecks in plasmid and virus generation (Fig. 3d), so further analysis was required to determine whether the viral library was sufficiently diverse.

We therefore computed how the fraction uniquely labeled neurons scaled with the diversity of the viral library. This problem is formally equivalent to a generalization of the classic “birthday problem”, which concerns the probability that, in a group of *k* people (neurons), some pair will have the same birthday (barcode). Assuming that all barcodes are equally abundant in the library, we can express the expected fraction *F* of uniquely labeled neurons as:

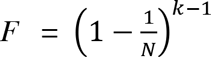
where *k* is the number of infected neurons (assuming one barcode per cell), and *N* is the barcode diversity (see Supplementary Note 2). Thus if *k*=1000 LC neurons were infected with a library of diversity of N=10^6^, on average 99.9% of all neurons would be labeled uniquely. This expression holds only for a library of equally abundant barcodes; if some barcodes are overrepresented the fraction *F* of uniquely labeled neurons decreases (in the same way that if birthdays tend to fall on a particular day the probability of finding a shared birthday in a group increases (Munford, 1977)). However, analysis of the actual viral barcode library determined by sequencing (Fig. 3e) revealed that in practice these deviations from uniformity had only a minor effect (Supplementary Fig. 2a). Thus, under our conditions, the vast majority (>99%) of neurons will be uniquely labeled, even taking into account the uneven barcode distribution in the viral library (Fig. 3f; Supplementary Note 2).

We also used a second, more empirical approach, to estimate the extent of degenerate labeling. Since we used the same viral library to infect neurons in different animals, barcode sequences found in more than one animal represent degeneracy. We therefore looked for overlap in the recovered barcodes from four independent injections of the same virus library. Out of the 992 unique barcodes that we recovered from traced neurons, only three barcodes were present in more than one animal, and no barcode was present in more than two animals. This empirically measured rate of degenerate labeling is in close agreement with our expectations based on the theoretical considerations above. Moreover, two of the three repeated barcodes were among the most abundant barcodes in the virus library (Supplementary Fig. 2b), and would thus be expected *a priori* to have the highest probability of double labeling.This analysis provides an independent confirmation that the error rate due to non-unique labeling by the barcode library is very low in our experiments.

In addition to non-unique barcode labeling, MAPseq is subject to other errors that differ from those associated with conventional tracing approaches. We used several approaches to quantify these errors, and find that the overall MAPseq error rate was low both for false positives (1.4+/
0. 8% (mean+/-std error)) and false negatives (8.6+/-6%) (*see* Supplementary Note 3). MAPseq thus provides a reliable measure of axon projections.

### Sequencing of barcode mRNAs reveals diverse single neuron projection patterns

The goal of MAPseq is to quantify the projection patterns of large populations of neurons in parallel. We therefore developed a method to determine the amount of each barcode in each dissected target (Fig. 4a). Forty-four hours after injection of barcoded virus into right LC, we performed reverse transcription on barcode mRNA extracted from dissected target regions. To overcome distortions introduced during amplification (Kebschull and Zador, 2015), and to allow a precise count of barcode cDNA molecules, we designed reverse transcription primers to tag each individual barcode mRNA molecule with a random 12-nt unique molecular identifier (UMI). We also added a 6-nt slice-specific identifier (SSI), to allow multiplexing of samples within a single high-throughput sequencing flow cell. We then amplified, pooled and sequenced these SSI-UMI-barcode cDNAs (Supplementary Fig. 3). We developed a conservative computational pipeline to minimize noise due to RNA contamination and to correct for sequencing and other errors (Supplementary Note 4). Finally, we converted barcode abundance in the target areas to a matrix of single neuron projection patterns.

**Figure 4:**
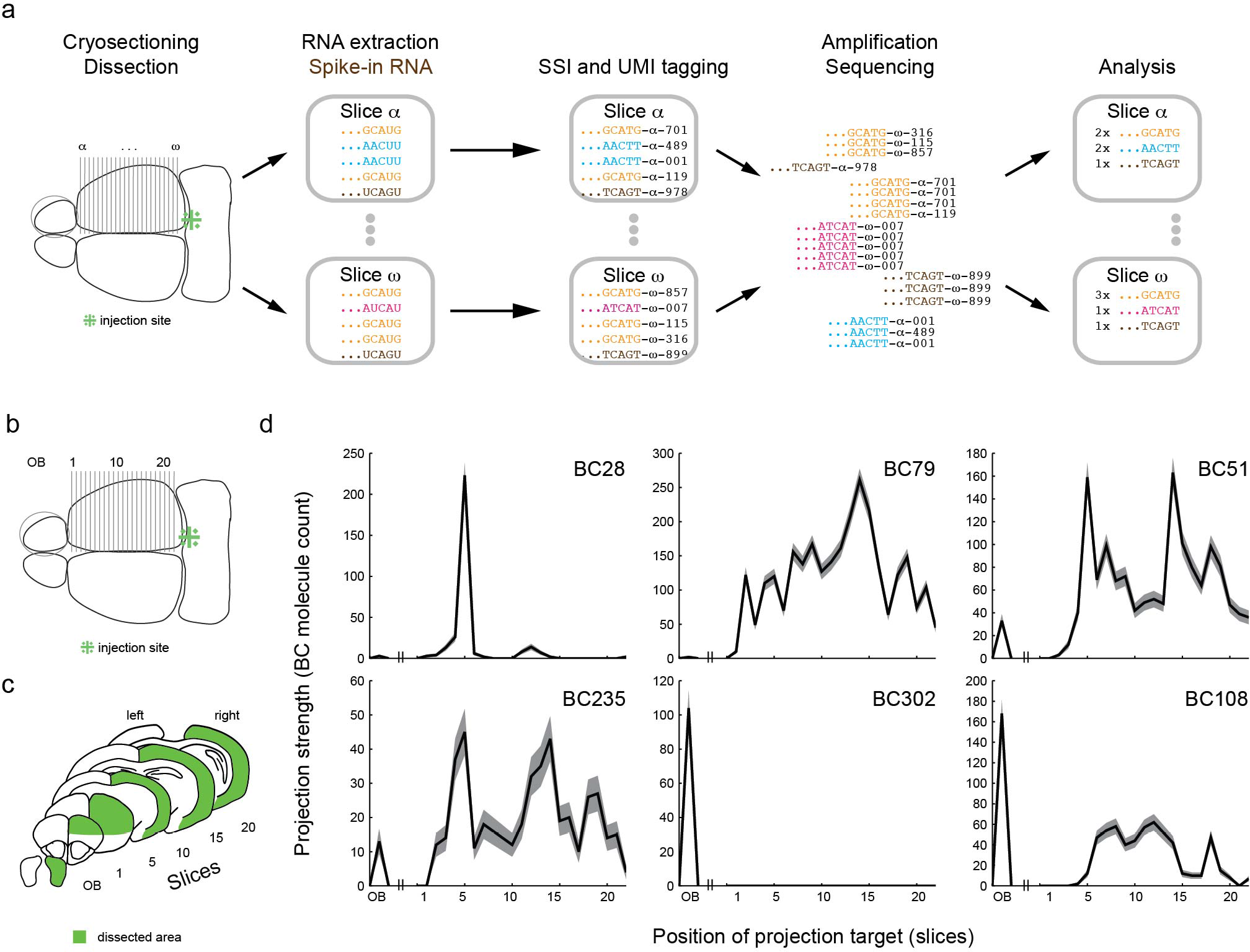
MAPseq reveals large diversity of projections from LC. (a) Barcode mRNAs from target areas are sequenced as described (SSI=slice specific identifier, UMI=unique molecular identifier). (b,c) Barcodes from ipsilateral olfactory bulb and cortex show projection patterns (d)with single or multiple peaks in cortex and/or olfactory bulb. The shaded area indicates Poisson error bars given by the square root of barcode (BC)counts perslice.

We used MAPseq to determine the projection patterns of a total of 995 barcodes labeled in four animals (249+/-103 barcodes per animal), roughly corresponding to an equal number of LC neurons. For each animal, we analyzed the barcode mRNAs extracted and amplified from the olfactory bulb and from 22 coronal slices (300μm) taken from the cortex ipsilateral to the LC injection (Fig. 4b,c).Although like conventional GFP tracing, MAPseq does not distinguish between synaptic connections and fibers of passage, we minimized the contribution of large fiber tracts in the white matter during dissection, so most of the mRNA barcode signal was likely generated from axons terminating in the regions of interest. Dissection of coronal slices allowed us to probe the organization of projections along the rostral-caudal axis, but we were insensitive to any additional structure along the medio-lateral axis. Because individual barcode mRNA molecules are tagged with a UMI before amplification, we obtained a precise quantification (subject to Poisson counting statistics; see Supplementary Note 5) of the amount of each barcode sequence in each dissected target. In this way we could infer the projection strength—the density of axon per tissue volume—of each neuron to each coronal target area. For example, we recovered 223 copies of BC28 in slice 5, but none in slice 20, indicating that the projection strength to slice 5 is at least a factor of 200 higher than our detection floor (Fig. 4d).

Inspection of the projection patterns revealed that, in contrast to the simplest prediction from conventional bulk tracing, single neurons did not project uniformly throughout the ipsilateral hemisphere. Instead, neurons projected in diverse and idiosyncratic ways to specific targets, innervating some areas hundreds of times more strongly than others (Fig. 4d). Some neurons (e.g. BC28) showed specific projections to only a small part of cortex, whereas others (e.g. BC79) projected more broadly, or projected to multiple areas (e.g. BC51 or BC235). Projections to the olfactory bulb appeared independent of projections to cortex, with some neurons projecting exclusively to the olfactory bulb (e.g. BC302), some projecting exclusively to cortex (e.g. BC79) and others projecting to both (e.g. BC108).

The small fraction of multiply infected neurons revealed by single cellsequencing (21+/-11%; Fig. 3b) provided a convenient internal measure of the reliability of MAPseq. Within each animal, we found pairs of very similar projection patterns, as would be expected if they arose from double labeling of the same neuron (Fig. 5a; Supplementary Fig. 4a). By comparing the similarity of projection patterns within and across animals, we estimated the total number of barcode pairs arising from doubly labeled neurons (18.6+/-2%; Fig. 5b; Supplementary Fig. 4b). The fact that this estimate is in such close agreement with the independent estimate of the number of doubly labeled neurons from single cell sequencing supports the view that MAPseq provides a robust measure of single neuron projection profiles.

**Figure 5:**
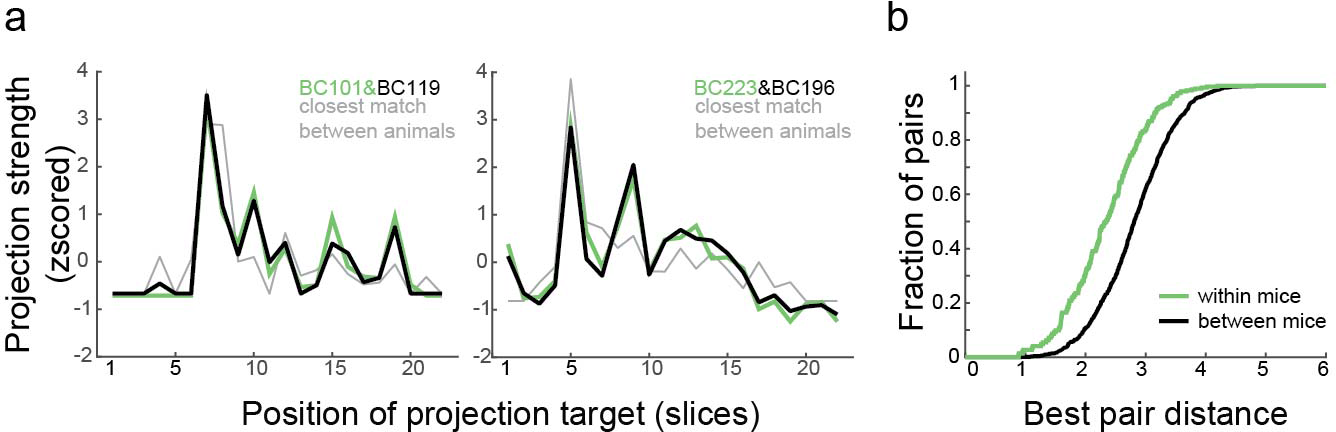
MAPseq provides a robust readout of single neuron projection patterns, (a) Two representative pairs of barcodes with projection patterns more similar than expected by chance for two distinct neurons, likely the result of double infection of a single neuron. The close agreement between the two barcode profiles indicates that MAPseq provides a reliable measure of projection patterns. The closest match across animals is indicated in rey for comparison, (b) Cumulative distribution of distances between the best barcode pairs within one animal and across animals. The shift in the within animal distribution reflects the higher fraction of closely matched projection profiles, consistent with double infection. Representative data from one animal.

To assess the structure of the LC projection to cortex and olfactory bulb, we sorted all traced neurons by their maximum projection (Fig. 6a). The maximum projections of individual LC neurons tile the entire cortex. To compare across the population we normalized the total barcode content of each neuron to one, although interestingly there was no correlation between expression level in the LC and the maximal projection strength to cortex, as would have been expected if differences across neurons were dominated by expression level (*R*=-0.06; p=0.09; Supplementary Fig. 5a). Only in the aggregate do these projections re-create the apparently homogeneous LC innervation of cortex previously described by bulk methods (Fig. 2e). Consistent with previous results (Shipley et al., 1985), a considerable fraction (23+/-4.7%) of all mapped neurons projected to the olfactory bulb.

**Figure 6:**
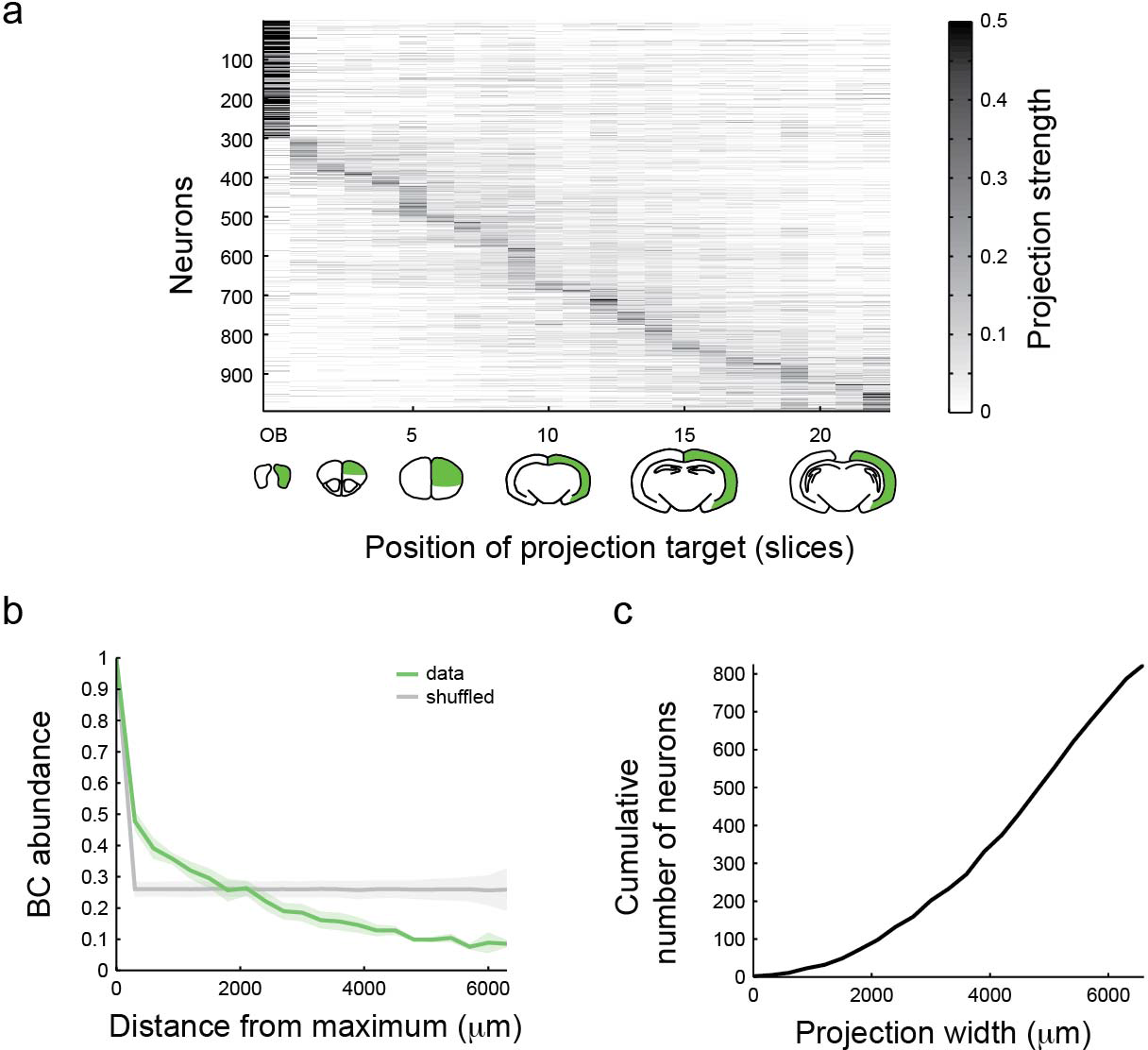
LC neurons tile cortex with their maximum projections, but innervate large areas of cortex at a low level, (a) A heatmap of all 995 projection patterns from 4 animals shows a strong diagonal component after sorting by maximum projection site. Barcode abundances are normalized to sum to one across target areas and are color-coded as indicated, (b) Average cortical drop-off rate from maximum for all barcodes shows a rapid drop-off and a structure that is different from the drop-off after randomly shuffling slices for all neurons. N=4. (c) Cumulative distribution of cortical projection widths indicates a broad low intensity innervation of cortex by individual LC neurons. BC=barcode.

We then asked if we could find structure, or even stereotypic projection cell types, in the single cell data set. We investigated the cortex-wide projection patterns of LC by reducing the dimensionality of the projection dataset using Euclidean distance based t-SNE (t-distributed stochastic neighbor embedding) (Van der Maaten and Hinton, 2008). Neurons with maximum projections close to each other in physical space along the rostro-caudal axis also fall closely together in t-SNE space (Supplementary Fig. 5b), indicating that the location of the maximum projection target at least partially describes the individual neuron projection patterns. However, hierarchical clustering of the projection profiles of neurons that project to cortex did not uncover distinct cell classes (Supplementary Fig. 5c). Although we cannot rule out the possibility that there is further structure in the projection patterns that would be revealed by higher-resolution dissection, our data suggest the intriguing possibility that LC projections are equipotential for projecting to all targets, and the choice is arbitrary for each neuron. How the circuit might exploit such random connectivity raises interesting computational challenges.

Although many LC axons projected very strongly to a narrow target, these axons often sent minor collaterals to a much broader area, like a plant with a single main stalk and many minor growths. The average number of projection peaks per LC neuron was 1.6+/-0.8 (Supplementary Fig. 5d), and the fall-off to half the maximum projection strength of individual neurons occurred on average in <300μm (Fig. 6b). Nonetheless, every cortically-projecting neuron innervated on average 65+/-23% of cortex at a detectable level (see cumulative distribution of projection widths in Fig. 6c). Importantly, this broad, weak innervation of cortex cannot be explained simply by contamination of our dataset by fibers of passage. LC axons innervate cortex starting in the rostral end and moving caudally (Fig. 3c). We would therefore expect signals from accidentally dissected fibers of passage to only precede strong projection targets of individual neurons along the rostro-caudal axis. However, we find that strong projection targets are both preceded and followed by low level projections (see e.g. Fig 4d BC79 and BC51). We therefore conclude that the observed weaker signals are not the result of fibers of passage but represent real, but weak, projections.
The fact that many neurons had a strong preferred target in cortex or olfactory bulb, but also projected weakly to a much broader area, provides a way to reconcile apparently conflicting results about the specificity of LC projections. Recent experiments, in which retrograde viral labeling was combined with anterograde tracing of axons, concluded that as a population LC neurons project largely indiscriminately throughout cortex and the rest of the brain examined (Schwarz et al., 2015). However, using this approach, a neuron labeled retrogradely from a weak projection is indistinguishable from one labeled from a strong projection, so at the level of the population (i.e. after summing the projection patterns of strongly and weakly projecting neurons) it may appear that projections are nonspecific. Thus although the results of this study may appear to contradict those obtained by MAPseq at single neuron resolution, simulations of retrograde labeling in combination with anterograde bulk tracing based on our MAPseq data set demonstrate that there is no contradiction (Supplementary Fig. 5e,f) at the bulk level.

### MAPseq scales to several injection sites

MAPseq can readily be extended to determine the projections of two or more regions in a single animal. As a proof-of-principle, we injected the same library of MAPseq virus bilaterally into left and right LC, and dissected coronal slices of left and right cortex and the olfactory bulbs, as well as both injection sites (Fig. 7a). Each barcode was expressed predominantly in either the left or right LC (Fig. 7b); barcode expression at the site contralateral to the injection, due to contralaterally projecting fibers and/or contamination, is much lower. Thus each barcode can be reliably assigned to the appropriate injection site. As expected, parallel injections recapitulated the projection pattern observed with single injections (Fig. 7c). Multiplexing MAPseq to dozens of injections per animal may be feasible, reducing the labor and cost of brain-wide projection mapping efforts, and eliminating the need to map data from multiple animals to an average reference brain (Oh et al., 2014; Zingg et al., 2014).

**Figure 7:**
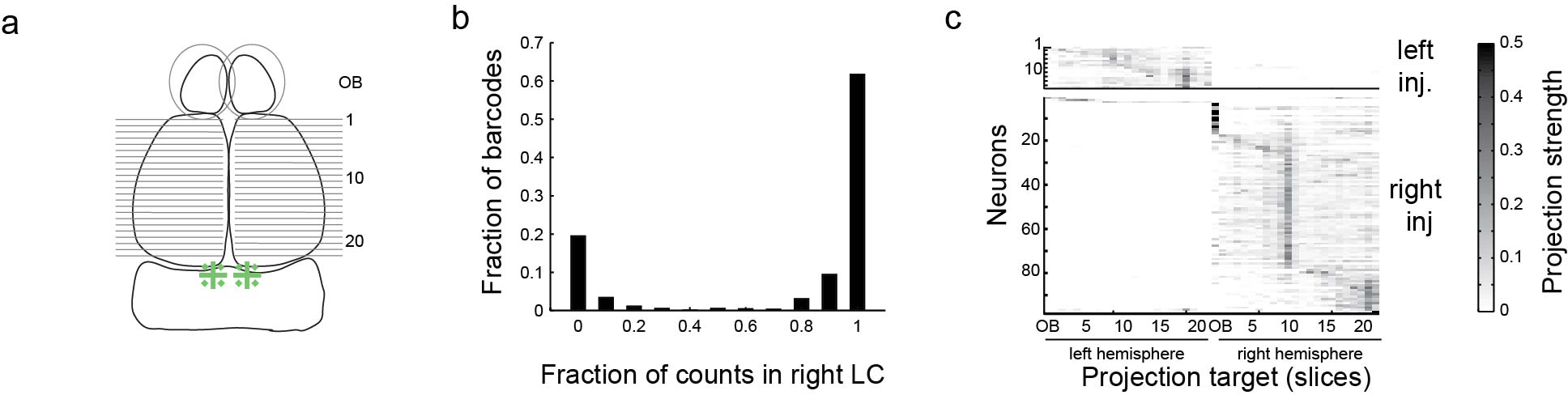
MAPseq can be multiplexed to several injection sites, (a) Following bilateral injection of barcoded Sindbis virus into LC, left and right olfactory bulb and cortex were dissected as before, (b) Histogram of the fraction of barcode counts in the right vs. left injection site across barcodes. Barcodes show strong abundance differences in the left and right injection sites allowing them to be assigned to one of the two sites, (c) Bilateral injections produce the projection pattern expected from unilateral injections. Differences in the number of neurons traced from the left and right LC arise from injection variability.

## Discussion

We have described MAPseq, the first application of sequencing to neuroanatomical tracing. MAPseq is a simple, rapid, and inexpensive approach to determining the projection patterns of myriad single neurons at one or more injection sites in a single animal. As a proof-of-principle, we applied MAPseq to the LC. In contrast to previous bulk labeling studies that reported diffuse and non-specific projections from the LC, this single neuron resolution analysis reveals that individual LC neurons have idiosyncratic projection patterns with preferred cortical targets, and reconciles a controversy about the specificity of LC projection patterns. MAPseq, which complements rather than replaces conventional approaches, can readily be applied to other brain regions and organisms, and with further development may be combined with information about gene expression and neural activity.

### High-throughput sequencing and neuroanatomy

The cost of sequencing the human genome was several billion dollars in 2003, but today it is less than one thousand dollars—a decrease of over six orders of magnitude in just over a dozen years (Hayden, 2014; Sheridan, 2014). This precipitous drop in sequencing costs continues unabated, at a rate faster even than Moore’s law (the rate at which computers improve). At the same time, DNA sequencing has evolved from a specialized tool for determining the sequences of genomes into a versatile technology for determining gene expression levels, discovering new species, tracking cell fates, and understanding cancer growth, among many other applications (Reuter et al., 2015). With advances of technology and novel assays, DNA sequencing has revolutionized disparate areas of biology. By harnessing sequencing for neuroanatomical tracing, MAPseq may accelerate our understanding of neural circuits.

Information about single neuron projection patterns obtained by MAPseq may be combined with information about other dimensions of neuronal function. For example, single neuron projection patterns obtained by MAPseq could be associated with information about gene expression. One approach exploits transgenic mice that mark defined neuron classes with Cre recombinase.Although expression of barcodes delivered using the RNA virus Sindbis cannot readily be controlled with Cre, barcodes could be delivered using a DNA virus like AAV or a retrovirus like lentivirus. More general approaches, such as single cell isolation (Supplementary Fig. 6), might associate several genes or even a whole transcriptome with projection patterns. MAPseq data could also be combined with recordings of neural activity obtained by calcium imaging. Taken together, the combination of connectivity, gene expression, and activity could provide aricher picture of neuronal function than any of these alone (Marblestone et al., 2014).

### MAPseq resolution

The factors limiting MAPseq can be understood by analogy with those limiting optical microscopy. Just as spatial resolution — effective pixel size — in microscopy is limited by the optics and detectors, sothe spatial resolution of MAPseq is limited by the spatial precision with which brain regions are defined. In the present work we used gross dissection, which affords only relatively crude spatial resolution, perhaps to the level of cortical areas (we have successfully dissected individual cortical areas, including the primary auditory cortex; Supplementary Fig. 7a,b). The spatial resolution can be increased by other methods including laser-capture microdissection (Espina et al., 2006), transcriptome in vivo analysis (TIVA) tagging (Lovatt et al., 2014), or fluorescent *in situ* sequencing (Lee et al., 2014) to determine the location of barcode mRNAs in tissue.

The effective spatial resolution of MAPseq may also be limited by the amount of barcode mRNA in each spatially defined region, analogous to the low-light photon-counting limit in microscopy. The precise spatial scale at which the barcode mRNA “shot noise” limit is reached in MAPseq is determined by the interplay of several factors, including the diameter of the axonal projections, expression level of the barcode, and the efficiency with which barcodes can be recovered and amplified. Because mRNA shot noise did not appear limiting in the present experiments, we did not invest significant effort in optimizing these parameters. However, the fact that barcodes can be detected even in fine LC axons suggests that MAPseq may achieve relatively high spatial resolution.

### Uniformity vs. specificity of LC projections

The LC sends projections to most ipsilateral brain areas, with the notable exception of the striatum. However, how broadly individual neurons innervate those target areas is subject to debate. Classical retrograde tracing studies suggested a topographic organization of neocortical (Waterhouse et al., 1983) and brain-wide (Loughlin et al., 1986) projection neurons in LC. Consistent with this, double retrograde labeling studies reported that the LC projections to frontal and motor cortices (Chandler et al., 2014; Chandler and Waterhouse, 2012) overlap minimally. In contrast, other double retrograde studies found overlap between neurons projecting to separate structures along the same processing stream (Simpson et al., 1997), or structures as different as forebrain and cerebellum (Steindler, 1981). More recent work using retrograde viral labeling combined with anterograde tracing concluded that LC neurons project largely indiscriminately throughout both cortex and the rest of the brain (Schwarz et al., 2015).

Our single cell resolution data reconcile these conflicting datasets. We find that individual LC neurons have very specific projection targets in cortex and olfactory bulb, but are not limited to a single target. We further observe that many LC neurons that project to cortex innervate a large fraction of the cortex at least weakly, in addition to having preferred projection targets. To the extent that retrograde viral tracers may not distinguish between strong and weak projections, it may be this feature of single neuron projections in combination with bulk tracing that lead Schwarz and colleagues (Schwarz et al., 2015) to conclude that LC neurons largely indiscriminately project throughout cortex and the bulb, and indeed we can replicate their results by simulating retrograde tracing on our single cell dataset.

The LC is the sole source of noradrenaline to the cortex. Noradrenaline exerts a powerful influence on an animal’s behavioral state. Noradrenaline levels control the overall level of vigilance; they are lowest during sleep, and are increased in response to stimuli such as pain. Noradrenaline gates attention, enhances formation of long-term memory and is thought to regulate the exploration-exploitation balance (Aston-Jones and Cohen, 2005; Sara, 2009). Traditionally, it has been assumed that the levels of neuromodulators such as noradrenaline represent a global signal, broadcast indiscriminately throughout the cortex. However, the specificity of the single neuron projections patterns uncovered by MAPseq suggests that different brain regions could be subject to differential control. Whether this potential for differential control is actually realized, and what functional role it plays, remain to be determined.

### Conclusion

Applying MAPseq to LC, we discovered unexpected structure that could not have been resolved by previous methods lacking single neuron resolution. MAPseq also lays the foundation for using sequencing to decipher local neuron-to-neuron connectivity (Zador et al., 2012). Using DNA sequencing technology, experimenters have gained unprecedented insight into the heterogeneity of cell populations at the single cell level (Navin et al., 2012). By leveraging this sequencing technology, MAPseq empowers neuroscience researchers to efficiently do the same for populations of projection neurons examined at the single neuron level.

## Experimental procedures

### MAPseq

Forty-four hours after injection of MAPseq virus into LC, we flash froze the brain and cut it into 300μm coronal sections using a cryostat. We dissected the cortical regions on dry ice and extracted total RNA from each sample. We then preformed gene specific reverse transcription for the barcode mRNA, produced double-stranded cDNA and PCR amplified it to produce an Illumina sequencing library which we sequenced at paired end 36 on an Illumina NextSeq sequencing machine.

### Data analysis

We processed the sequencing data to determine the exact copy number of each barcode sequence in each target area and in the injection site, an produced a barcode matrix, where each row corresponds to one specific barcode sequence, each column corresponds to a target area or the injection site and each entry is the copy number of that barcode mRNA in the respective area. All data are expressed as mean +/- standard deviation unless otherwise stated.

For full details on the experimental procedures please refer to Extended Experimental Procedure section in the supplement.

## Author contributions

I. D.P and A.M.Z conceived the study. J.M.K. and P.G.S. performed the experiments. J.M.K and A.P.R. performed the single cell isolation. J.M.K. and A.M.Z. analyzed the data. J.M.K. and A.M.Z. wrote the paper. A.M.Z. andD.F.A. supervised the project.

## Acknowledgements

The authors would like to acknowledge Vasily Vagin, Fred Marbach, Alex Koulakov, and Brittany Cazakoff for useful discussions and assistance, and Barry Burbach and Diana Gizatulliana for technical support. We would further like to thank Anirban Paul for help with single cell isolations. This work was supported by the following funding sources: National Institutes of Health [5RO1NS073129 to A.M.Z., 5RO1DA036913 to A.M.Z.]; Brain Research Foundation [BRF-SIA-2014-03 to A.M.Z.]; Simons Foundation [382793/SIMONS to A.M.Z.]; PhD fellowship from the Boehringer Ingelheim Fonds to J.M.K.; PhD fellowship from the Genentech Foundation to J.M.K.; PhD fellowship fromthe Fundagao para a Ciencia e Tecnologia, Portugal to P.G.S.; Pew Scholarship and CSHL startup funds to D.F.A. This work was performed with assistance from CSHL Shared Resources, which are funded, in part, by the Cancer Center Support Grant 5P30CA045508.

### Data deposition

All high throughput sequencing datasets are publicly available under SRA accession codes SRS1204613 (library ZL067; virus library), SRS1204589 (libraries ZL068, ZL070, ZL071, ZL072; unilateral MAPseq datasets), SRS1204614 (libraries ZL073, ZL074; bilateral MAPseq datasets), and SRS1204626 (libraries ZL075 and ZL078; single cell dataset).

### Conflict of interests

The authors declare no conflict of interests.

## Extended Experimental Procedures

### MAPP-nX

MAPP-nλ is a modified version of pre-mGRASP (Kim et al., 2011). We stripped the pre-mGRASP protein of the 2A-cerulean fusion and added four repeats of the nλ RNA binding domain (Daigle and Ellenberg, 2007)in the cytoplasmic tail after amino acid 287 of the original pre-mGRASP sequence. We also added a Myc epitope tag followed by the CLIP-tag domain (Gautier et al., 2008) after amino acid 59 of the original pre-mGRASP protein.

### Sindbis virus barcode library

The virus used in this study is based on a dual promoter pSinEGdsp construct (Kawamura et al., 2003). We inserted MAPP-nλ after the first subgenomic promoter. Downstream of the second subgenomic promoter, we inserted the GFP coding region followed by closely spaced NotI and MluI restriction sites and four repeats of the boxB motif (Daigle and Ellenberg, 2007). Using this construct, we produced a high diversity plasmid library by inserting a diverse pool of double stranded ultramers (Integrated DNA Technologies) with sequence 5’-AAG TAA ACG CGT AAT GAT ACG GCG ACC ACC GAG ATC TAC ACT CTT TCC CTA CAC GAC GCT CTT CCG ATC TNN NNN NNN NNN NNN NNNNNN NNN NNN NNN NYY GTA CTG CGG CCG CTA CCT A-3’ between the NotI and MluI sites. We then produced Sindbis virus as previously described (Kebschull et al., 2015) using either the conventional DH(26S)5’SIN helper (Bredenbeek et al., 1993) or the new DH-BB(5’SIN;TE12) (Kebschull et al., 2015) helper. We determined the titer of the resulting virus by qPCR as previously described (Kebschull et al., 2015) and determined the viral library diversity by Illumina sequencing of the RNaseI protected genomic virus RNA.

### Injections

Animal procedures were approved by the Cold Spring Harbor Laboratory Animal Care and Use Committee and carried out in accordance with National Institutes of Health standards.

We pressure injected 180nl of 2×10^10^ GC/ml barcoded Sindbis virus uni-or bilaterally into LC of 8-10 week old C57BL/6 males (Jackson Labs) as described (Cetin et al., 2007). We leveled the animal skulls on two axes using lambda and bregma for the AP axis and 2mm laterally from the midpoint between lambda and bregma for the lateral axis. We used coordinates AP=-5.4mm, ML=0.8mm, DV=2.9mm and 3.1mm forLC and measured depth from the surface of the brain. We injected each DV coordinate with 90nl of virus, waiting ten minutes in between each depth. We sacrificed animals 44 hours post injection. For immunofluorescence, RNA *in situ* and histology, we transcardially perfused animals with ice cold saline (9g/l) followed by 4% paraformaldehyde (Electron Microscopy Sciences) in 0.1M Phosphate buffer. For RNA work we extracted the fresh brain and flash froze it on dry ice.

For measurements of MAPseq efficiency, we injected red retrobeads (Lumafluor) into the right olfactory bulb of 8-12 week old C57BL/6 males (Jackson Labs). Briefly, we roughly determined the center of the right olfactorybulb, and measured +/-1mm from the center in the AP axis and performed two craniotomies 2mm apart. We sonicated the beads for 20 minutes prior to injection to homogenize the solution and injected 210nl of stock concentration of beads at three different depths (0.3mm, 0.6mm and 0.9mm DV from the surface of the olfactory bulb) as described (Cetin et al., 2007). Twenty-four hours later, we injected barcoded Sindbis virus into right LC as above and sacrificed the animals 44-48 hours after Sindbis injection.
Immunofluorescence and ISH

We performed anti-GFP staining and RNA *in situ* hybridization on 6μm thick paraffin sections. For immunofluorescence, we used a rabbit anti-GFP antibody ab290 (Abcam) after heat induced antigen retrieval. We performed RNA *in situ* hybridization using the Panomics ViewRNA ISH Tissue kit (Affymetrix) using anti-GFP probe VF1-10141 according to the manufacturer’s protocol (10 minutes boiling and 10 minutes protease treatment). We performed anti-TH staining on floating 70μm vibratome sections using rabbit anti-TH antibody SAB4300675 (Sigma-Aldrich).

### Spike-in RNA

To produce spike-in RNA, we double stranded an ultramer (Integrated DNA Technologies) with sequence 5’-GTC ATG ATC ATA ATA CGA CTC ACT ATA GGG GAC GAG CTG TAC AAG TAA ACG CGT AAT GAT ACG GCG ACC ACC GAG ATC TAC ACT CTT TCC CTA CAC GAC GCT CTT CCG ATC TNN NNN NNN NNN NNN NNN NNN NNN NAT CAG TCA TCG GAG CGG CCG CTA CCT AAT TGC CGT CGT GAG GTA CGA CCA CCG CTA GCT GTA CA-3’ and then *in vitro* transcribed the resulting dsDNA using the mMessage mMachine T7 *in vitro* transcription kit (Thermo Fisher) according to the manufacturer’s instructions.

### qPCR

We reverse transcribed total RNA using oligodT primers and Superscript III reverse transcriptase (Thermo Fisher) according to the manufacturer's instructions. We then quantified the amount of barcode and β-actin cDNA by qPCR in SYBR green power master mix (Thermo Fisher) according to the manufacturer's instructions using primers 5’-GAC GAC GGC AAC TAC AAG AC-3’ and 5’-TAG TTG TAC TCC AGC TTG TGC-3’ for barcode cDNA and 5’-CGG TTC CGA TGC CCT GAG GCT CTT-3’ and 5’-CGT CAC ACT TCA TGA TGG AAT TGA-3’ for β-actin cDNA.

### MAPseq

We cut 300μm thick coronal sections of fresh frozen brains using a Leica CM 3050S cryostat at −12°C chamber temperature and −10°C object temperature. To avoid cross-contamination between samples, we took care to cut each section with a fresh, unused part of the blade. We melted each section onto a clean microscope slide and rapidly froze the section again on dry ice before dissecting out the cortex on dry ice using acold scalpel blade. During dissection, we aimed to avoid known fiber tracts to minimize the contamination of our dataset with fibers of passage. After sample collection, we processed all samples out of order to avoid potential sample cross-contamination from impacting interpretation of MAPseq results.

We extracted total RNA from tissue samples using Trizol reagent (ThermoFisher) according to the manufacturer's instructions. We mixed the total RNA from the tissue samples with spike-in RNA. We then produced ds cDNA as previously described (Morris et al., 2011) using a gene specific primer of from 5’-CTT GGC ACC CGA GAA TTC CAN NNN NNN NNN NNX XXX XXTGTA CAG CTA GCG GTG GTC G-3’, where XXXXXX is one of 65 trueseq like SSI and N_12_ is the UMI. We then cleaned the reaction using the Qiagen MinElute PCR purification kit according to the manufacturer’s instructions and treated the eluted ds cDNA with ExonucleaseI (New England Biolabs) to remove remaining primers. We amplified the barcode amplicons by nested PCR using primers 5’-CTC GGC ATG GAC GAG CTG TA-3’ and 5’-CAA GCA GAA GAC GGC ATA CGA GAT CGT GAT GTG ACT GGA GTT CCT TGG CAC CC GAG AAT TCC A-3’ for the first PCR and primers 5’-AAT GAT ACG GCG ACC ACC GA-3’ and 5’-CAA GCA GAA GAC GGC ATA CGA-3’ for the second PCR in Accuprime Pfx Supermix (Thermo Fisher). We then gel extracted the amplicons using the Qiagen MinElute Gel extraction kit according to the manufacturer's instructions and pooled the individual sequencing libraries based on qPCR quantification using primers 5’-AAT GAT ACG GCG ACC ACC GA-3’ and 5’-CAA GCA GAA GAC GGC ATA CGA-3’. We then sequenced the pooled libraries on an Illumina NextSeq500 high output run at paired end 36 using the SBS3T sequencing primer for paired end 1 and the Illumina small RNA sequencing primer 2 for paired end 2.

### Efficiency measurements and single cell isolation

After transcardial perfusion with ice-cold artificial cerebrospinal fluid (127mM NaCl, 25mM NaHCO_3_, 1.25mM NaPO_4_, 2.5mM KCl, 2mM CaCl_2_, 1mM MgCl_2_, and 25mM D-glucose), we extracted the unfixed brain and flash froze the bead-injected olfactory bulb on dry ice before processing it for sequencing as described above. We cut 400μm thick acute sagittal slices of the remaining right hemisphere in dissection solution (110mM choline chloride, 11.6mM ascorbic acid, 3.1mM Na pyruvic acid, 25mM NaHCO_3_, 1.25mM NaPO_4_, 2.5mM KCl, 0.5mM CaCl_2_, 7mM MgCl_2_, and 25mM D-glucose) using a Microm HM650V vibratome. We incubated sections containing LC in artificial cerebrospinal fluid (126mM NaCl, 20mM NaHCO_3_, 3mM KCl, 1.25mM NaH_2_PO_4_, 2mM CaCl_2_, 2mM MgSO_4_, and 20mM D-glucose) containing synaptic blockers (0.05mM APV, 0.02mM DNQX and 0.1 μM TTX) for 20 minutes at room temperature. We then digested the slices in artificial cerebrospinal fluid with streptomyces griseus protease (Sigma P5147) at 1mg/ml at room temperature for 30 min. After washing in artificial cerebrospinal fluid with synaptic blockers, we dissected LC from the digested section and triturated the tissue to produce a single cell suspension. Using an inverted fluorescent microscope (Zeiss Observer), we picked individual cells by hand, deposited the cells directly into lysis buffer (2.4μl 0.2% triton, 1μl 10mM dNTPs, 1μl 10mM RT primer per cell) and proceeded to preparing sequencing libraries from the cells as described above for tissue samples.

Animals used.

Animals used
| Number of animals | Manipulation | Figures based on these animals |
| --- | --- | --- |
| 4 | Right LC injection with MAPseq virus; dissection of right cortex and olfactory bulb; qPCR and sequencing of barcode RNA | Fig. 2e, 4, 5, 6<br>Supp. Fig. 1, 2, 4, 5, 7c,d |
| 2 | Right LC injection with MAPseq virus; dissection of right and left cortex; qPCR of barcode RNA | Fig. 2f |
| 1 | Bilateral LC injection with MAPseq virus; dissection of right and left cortex and olfactory bulb; qPCR and sequencing of barcode mRNA | Fig. 7 |
| 3 | Right LC injection with MAPseq virus and retrobeads injection into right olfactory bulb; single cell isolation from LC | Fig. 3b;<br>Supp. Fig. 6 |
| 3 | Right LC injection with DH(26S)5'SIN packaged MAPseq virus and dissection of select cortical targets and the olfactory bulb and sequencing of barcode mRNA | Supp. Fig. 7a,b |
| 6 | Right LC injection with DH(26S)5'SIN packaged MAPseq virus. TH staining of LC and quantification of overlap and count of TH+ neurons | Supp. Fig. 1j,k,l |
| 3 | Right LC injection with DH(26S)5'SIN packaged MAPseq virus; ISH for barcode mRNA and IF for GFP protein | Fig. 3b<br>Supp. Fig. 1 |
| 2 | Right LC injection with DH(26S)5'SIN packaged MAPseq virus and dissection of the olfactory bulb and ipsilateral cortex and sequencing of barcode mRNA | Supp. Fig. 1e,f |

## Supplementary Notes

### Supplementary Note 1: Sindbis virus

MAPseq requires that a propagation-incompetent virus be used for barcode delivery, i.e. after a neuron is infected with a particular barcode, the virus carrying this barcode should not propagate and spread to other cells. If the virus did propagate, barcodes would spread from cell to cell, and unique labeling of neurons by barcodes would break down as many neurons would now share the same barcode.

Initial Sindbis virus libraries prepared with the conventional helper construct DH(26S)5’SIN induced GFP labeling not only at the injection site, but also occasionally at sites far away from the primary site of injection (Supplementary Fig. 1). Such distal labeling has previously been interpreted as retrograde infection (Furuta et al., 2001), but we recently showed that it does not arise from retrograde spread, but is due instead to secondary infection (Kebschull et al., 2015). Consistent with secondary spread, a subset of barcodes showed unexpectedly high expression levels in single target areas (“spikes”; Supplementary Fig. 1e) when we performed MAPseq with barcoded virus packaged using the conventional helper construct. Indeed, we find that the barcode expression level in these spikes is comparable to the expression level of the same barcodes at the primary injection site (Supplementary Fig. 1f). These observations strongly suggested that the observed spikes originate from ectopically infected cell bodies that are labeled with the same barcode as a neuron at the injection site. Given the high diversity of the viral library, such double labeling is exceedingly unlikely to occur by chance if labeling were due to retrograde infection, but would be expected if the virus used propagated inside the brain.

We therefore designed a new helper construct, (DH-BB(5’SIN;TE12ORF); (Kebschull et al., 2015)), to eliminate secondary spread. When we used this modified helper construct, which minimized co-packaging of Sindbis virus Defective Helper RNA, we almost completely eliminated secondary infection, and were unable to detect any more spikes by sequencing (Supplementary Fig. 1g). Sindbis virus packaged by DH-BB(5’SIN;TE12ORF) thus fulfills the requirements for use in MAPseq. In all subsequent MAPseq experiments we used viral libraries prepared with the modified helper virus.

### Supplementary Note 2: Labeling neurons with barcodes

In MAPseq we randomly label neurons with barcodes from a viral library to provide them with a unique identity. Ideally, every infected neuron would have a single, unique barcode. There are two deviations from this ideal scenario: (i) multiple neurons per barcode; and (ii) multiple barcodes per neuron. We consider the implications of the former in more detail below.

### Multiple neurons per barcode

Multiple neurons per barcode, i.e. degenerate labeling, is problematic as it leads to incorrect results. Consider for example two neurons, A and B, that project to distinct cortical target areas. If by chance A and B are labeled with the same barcode (e.g. barcode 11), then MAPseq will return the merged projection pattern of A and B as the projection pattern of barcode 11 (Fig. 3c). While this is indeed the projection pattern of barcode 11, it cannot be interpreted as the projection of a single neuron. Errors of this type can be avoided by using sufficiently diverse viral libraries, thereby minimizing the probability that the same barcode will label two different neurons. This implies that the requisite diversity of the viral library depends on the number of neurons infected.

Here we formulate the mathematical problem: Given a population of *k* neurons, labeled randomly from a pool of *N* barcodes, what is the probability that a given neuron will be uniquely labeled? This is closely related to the problem: What is the probability thata given barcode will appear in more than one neuron? These problems are related to the classical problem of drawing balls with replacement from an urn, where every ball corresponds to a barcode sequence, and the probability of drawing each ball is determined by the abundance of this barcode in the library.

We first consider a simplified case, in which we assume that every barcode is equally abundant in the virus library, i.e. that the barcode probability distribution is uniform. What then is the expected number of neurons that share a barcode with at least one other labeled cell? If there are only two neurons A and B, then the probability of neuron B having the same barcode as neuron A is P(A)=1/*N*, so the probability that A's barcode is unique is 1-P(A). Generalizing to *k* infected neurons, the probability that A's barcode is unique is (1-P(A))^(k-1)^, and the probability that it is not unique is 1-(1-P(A))^(k-1)^. As the expected value of a sum is the sum of its expected values, the expected number of non-uniquely labeled neurons is

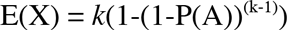

The fraction of uniquely labeled neurons F is then

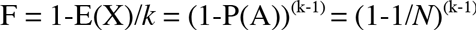

Similarly, the expected number *D* of barcodes used morethan once is 
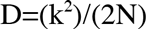
swhere *N* is the number of barcodes and *k* is the number of infected cells, and we have assumed N>>k.

Now, let us consider the more realistic case, in which the distribution of barcode abundance is not uniform, so neurons are more likely to be labeled with some barcodes than others. To calculate the expected value of non-uniquely labeled neurons in this case, we generalize the reasoning above by including a sum over all barcodes, weighted by their probability. E(X) is then given by

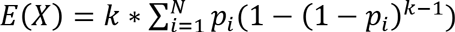
where *p_i_* is the probability of barcode i=1‥N, *k* is the number of infected neurons and *N* is the total number of barcodes in the virus library.

To determine the empirical distribution of barcodes in the virus library, we directly sequenced the genomic RNA of an aliquot of our Sindbis virus. Sequencing was performed at sufficient depth to overcome Poisson sampling introduced by Illumina sequencing. After error correction, the absolute abundance of different barcode sequences is a direct measure of the barcode probability distribution (Fig. 3e). Despite error correction, there is a chance of including erroneous barcode sequences when counting barcodes that have a very low molecule count.For all calculations, we therefore chose a conservative threshold, and required at least 3 counts for barcodes to be included in the virus library. Based on this empirically determined distribution of barcode abundances, we then calculated the fraction of uniquely labeled cells as a function of the number of infected cells based on the above derivations (Fig. 3f). Simulations indicate that removing the most abundant barcodes have little effect on the capacity of the library to label neurons uniquely (Supplementary Fig. 2a). These results indicate that the observed nonuniformity in the abundance of barcodes in the library does not substantially interfere with the capacity of the library to uniquely label large numbers of cells.

### Supplementary Note 3: False positive and negative rates of MAPseq

#### MAPseq false negative rate

Like every experimental method, MAPseq is susceptible to both false negatives and false positives. First, we sought to relate the efficiency of MAPseq and thus its false negative rate to established neuroanatomical methods. MAPseq is conceptually closest to GFP-based methods (Oh et al., 2014; Zingg et al., 2014), in which a genetically-encoded fluorophore is expressed in a neuronal population, and fluorescence is detected in targets. The sensitivity and selectivity of such fluorophore-based methods depend on many factors, including expression level, imaging conditions, background fluorescence, etc. To our knowledge there has not been a rigorous and precise quantification of the sensitivity and selectivity of such methods, which would allow us to compute e.g. the probability of detecting a small axon for e.g. a given fluorophore expression level, etc; nor indeed is it clear how one would ground-truth such a quantification. Moreover, direct comparison of MAPseq and fluorophore-based methods on a section-by-section basis would be challenging because the optimal conditions for imaging and RNA extraction differ. We therefore did not attempt a quantitative comparison of the efficiency of MAPseq with that of fluorophore-based methods.

Instead, we compared the efficiency of MAPseq to that of another well-established method, Lumafluor retrobeads which allows us to directly compare the efficiency of bead labeling and MAPseq within the same animal. Briefly, we injected red retrobeads into the olfactory bulb, and MAPseq Sindbis virus into LC (Supplementary Fig. 6). Retrobeads taken up by axons in the olfactory bulb are actively transported back to cell bodies and label bulb-projecting cells. Barcodes from infected LC cells that are labeled with retrobeads should therefore be present in the bulb. The fraction of barcodes recovered from retrobead-labeled LC neurons that are also detected in the olfactory bulb by MAPseq thus provides a neuron-by-neuron estimate of the MAPseq false-negative rate.

To calculate this measure of efficiency, we performed MAPseq on the olfactory bulb and sequenced the barcode complement of individual bead and Sindbis labeled LC cells by dissociating LC, and picking individual red and green cells using glass pipets (Sugino et al., 2006). Producing a single cell suspension from tissue slices involves digestion of the extracellular matrix and trituration of the tissue, which inevitably leads to breaking of processes and release of barcode mRNA into the bath. Given the very high expression levels of Sindbis virus, it was critical to determine the contribution of barcodes present freely floating in the bath or in cell debris, as these barcodes will be collected alongside the labeled cells and sequenced, and will later be indistinguishable from cell resident barcodes except for their abundance. We measured this background noise distribution by collecting cells that were GFP-negative, but were bead labeled. Since GFP-negative neurons do not express barcodes, any barcodes recovered from such cells represent contamination. We used the level of such contamination to establish the threshold for true barcode expression in intact isolated neurons.

We collected 45 neurons that were labeled with both GFP/barcodes and with red retrobeads from the olfactory bulb, and 9 neurons labeled only with red retrobeads to determine the background noise level of barcode expression. We found that MAPseq efficiency is high: 91.4+/-6% (mean +/- std error) of all barcodes from cells that project to the bulb as determined by bead labeling also appear to project to the bulb by sequencing (across 3 animals; Supplementary Fig. 6d). This estimate is robust over a large range of reasonable estimates for the level of background barcode contamination (Supplementary Fig. 6f). We therefore conclude that the false negative rate of MAPseq is 8.6+/-6%.

#### MAPseq false positive rate

A false positive event in MAPseq is the detection of a barcode in a target area to which the neuron expressing the barcode does not project. There are two potential sources of false positives. First, we might correctly detect a barcode that does indeed target this particular area, but we might mistakenly identify it as a different barcode (due e.g. to sequencing errors). Alternatively, barcodes that arise from other samples (slices), or from outside sources, might contaminate the target sample. (A third kind of error, those arising from insufficient barcode diversity, might also be considered a special case of false positives, but are considered separately above in “Unique labeling of neurons with barcodes”).

Due to the large combinatorial space of barcodes, it is exceedingly unlikely to mistake one barcode for another because of PCR or sequencing errors (see Supplementary Note 4). Contamination, however, is a concern and needs to be quantified.

LC neurons project primarily to the ipsilateral hemisphere (Waterhouse et al., 1983), and only a small fraction of LC neurons project to both ipsilateral and contralateral cortex (Room et al., 1981). Quantifying the projection strength of neurons to the contralateral hemisphere relative to their projection to the ipsilateral hemisphere therefore provides an upper bound on the rate of contamination, and thus on the false positive rate of MAPseq. Note that samples from the ipsi-and contralateral hemisphere were processed intermixed and out of order. Cross-contamination between samples from the ipsi and contralateral side should therefore be comparable to contamination between samples from the ipsilateral side only, and should be a good measure of overall contamination levels.

We used the MAPseq dataset of the bilaterally injected animal (Fig. 7) to calculate this upper bound to the false positive rate. Briefly, we calculated the ratio of the total number of barcode molecules detected in the contralateral hemisphere to the total number of barcode molecules detected in the ipsilateral hemisphere for all barcodes that projected more strongly to the ipsi-than contralateral side (n=115). The mean ratio, and thus upper bound to the MAPseq false positive rate is 1.4+/-0.8% (mean+/-std error). Note that our assumption that LC neurons project only ipsilaterally is conservative; violations of this assumption would increase the estimated falsepositive rate. Thus we conclude that MAPseq has a low false positive rate. These results indicate that MAPseq provides both sensitive and reliable mapping of long-range projection targets of a large number of neurons.

### Supplementary Note 4: Bioinformatics

Raw MAPseq data consist of two.fastq files containing Illumina sequencing results, where paired end 1 covers the barcode sequence, and paired end 2 covers the 12-nt UMI and the 6-nt SSI (Supplementary Fig. 1h and 3). To convert these sequencing data into projection maps, we first preprocessed the data in bash, before analyzing them in Matlab (Mathworks).

#### Preprocessing of sequencing data

Briefly, we stripped the fastq files of their quality information and trimmed the reads to the relevant length, then merged paired end 1 and 2 into a single file. Each line of this file corresponded to a single read containing the 30-nt barcode, the 2-nt pyrimidine anchor (YY), the 12-nt UMI and the 6-nt SSI. We de-multiplexed the reads based on the SSI using the *fastx_barcode_splitter* tool and filtered the reads to remove any ambiguous bases. We then collapsed the reads to unique sequences and sorted them.

Next, we selected a threshold of how many reads a sequence has to have to be considered for analysis. We were guided by earlier work on the effect of PCR amplification during Illumina library generation on next generation sequencing data (Kebschull and Zador, 2015). In this previous work, we found that when amplifying a pool of unique barcode sequences by PCR, the sequence rank profile of the Illumina results consists of a plateau of sequences with roughly equal read counts, followed by a shoulder and a long tail. The tail of this distribution is formed almost exclusively by PCR errors. In the MAPseq datasets, we therefore manually selected a minimum read threshold to remove the tail of the sequence rank profile from the analysis. This avoids contamination of our dataset with large numbers of PCR and sequencing errors and simplifies subsequent error correction and analysis steps.

We then collapsed the remaining reads (30-nt barcode+YY+12-nt UMI) after removal of the 12-nt UMI to convert reads into molecule counts. Note that we here ignored any potential PCR or sequencing errors in the 12-nt UMI, which will lead to a slight, but uniform, overestimation of molecule counts as two copies of the same cDNA with an error in the UMI only will be counted as two distinct molecules rather than one.

#### Split of barcodes from spike-ins

Spike-in molecules are barcodes of length 24 followed by the constant sequence ATCAGTCA, and are therefore easily distinguished from barcodes expressed from the virus (Supplementary Fig. 1h). As they carry different information, we split the uncorrected barcode data into spike-ins (perfect match to N_24_ATCAGTCA) and virally expressed barcodes (no N_24_ATCAGTCA sequence, but N_30_YY) and processed them separately.

#### Error correction

A random barcode of 30nt length has a potential diversity of 4^30^≈10^18^ different sequences. If we sample a relatively small number of barcodes from this enormous diversity, the chosen barcodes are likely very different from each other. Therefore many mutations to any given barcode are necessary to convert it into any other barcode of the chosen set.

We exploited this fact to correct errors in the sequenced barcodes. Using the short read aligner *bowtie* (Langmead et al., 2009), we performed an all-against-all mapping of all barcode sequences with >1 counts, allowing up to 3 mismatches, and forcing bowtie to output all possible alignments. We then constructed a connectivity matrix of all barcode sequences, where bowtie alignments are the connections between sequences. We used Matlab to find all connected graph components, that is all barcodes that mapped to each other, and collapsed the molecule counts of each of the members of such a connected component to the sequence of the most abundant member. We then removed low complexity sequences—a common artifact of Illumina sequencing—by filtering barcodes with stretches of more than 6 identical nucleotides. Finally, we compared all error corrected barcode sequences to the error corrected barcode sequences foundin the original virus library and kept only those barcodes for analysis that had a perfect match in the virus library.

Code for preprocessing of all MAPseq libraries can be found in *preprocessing.sh* and *matlab_preprocessing.m.* The viral library was processed using *viruslibrary_preprocessing.sh* and *viruslibrary_matlabcode.m.*

#### Analyzing the projection pattern

The described workflow results in a list of barcode sequences and their molecule counts in each target area and the injection site. Using Matlab, we then matched the barcode sequences in the injection site (reference barcodes) with the barcode sequences in the target sites, constructing a barcode matrix of size [# of reference barcodes]x[# of target sites#+# of injection sites] which then acts as the basis of all further analysis. Note that barcodes that appear in target areas and not the injection site (‘orphans’) are very rare and have low abundances, consistent with an interpretation of orphan barcodes as contaminants.

To exclude low confidence projection patterns from analysis, we required each barcode to have more than 100 counts in the injection site and at least one target area with more than 30 counts.

Code can be found in *producebarcodematrix_unilateral.m* and *producebarcodematrix_bilateral.m.*

#### Barcode matrix normalization

Raw barcode counts are very useful to survey the data available and to form intuitions about the mapped projection patterns. However, to compute summary statistics, we normalized the raw barcode matrix. We first normalized each target area by the number of unique spike-in molecules detected in each, to normalize for varying reverse transcription, PCR or library making efficiencies. We then normalized each area by the amount of β-actin per μl of total RNA (as measured by qPCR) to correct for varying tissue input and RNA extraction efficiencies. Lastly, we normalized all barcodes to sum to 1 across all target areas to correct for different expression levels of different barcodes.

Code for all analysis of the barcode matrix can be found in *analyse_unilateralinjections.m* and *analyse_bilateralinjections.m.*

#### Peak finding

To summarize LC projection patterns, we set a number of criteria to define peaks for each barcode. First, peaks need be at least half as high as the maximal barcode count across all target sites. Second, peaks need to be separated by at least 3 slices, and third, peaks need to rise at least their half maximal height from their surroundings (‘prominence’). Code used to find peaks can the found in *detectpeaks.m.*

#### Identification of double-infected cells

In order to identify pairs of barcodes that originated from double-infected cells, we looked for projection profiles from individual mice that are more similar than expected for barcodes from different cells. Briefly, we calculated the minimum pairwise Euclidean distance of every barcode profile to any other barcode profile of a particular mouse in z-scored space (“within mouse”). We then constructed a null-distribution by repeatedly calculating the minimum pairwise distances for every barcode profile from that mouse to a random sample of the same size of barcodes profiles obtained by sampling the other three mice in this MAPseq dataset (“between mice”). Distances that appear in the “between mice” null-distribution result from the similarity of the projection profiles of different cells. Therefore, distances in the “within mouse” set lower than those explained by this null distribution suggest that the two barcode profiles are more similar than would be expected for two separate cells. The two barcodes that correspond to this low distance probably arise from a single double-infection cell. Accordingly, we defined those barcode pairs as originating from double-infected cells that had distances in the left tail of the null distribution subject to Bonferroni correction for multiple hypothesis testing.

To estimate the overall number of barcodes from double-infected cells in every animal, we calculated area between the probability density function of the “within mouse” distances and the “between mouse” distances and multiplied it by the total number of barcode pairs in the dataset. We then took the number of barcode pairs as our estimate of the number of barcodes in double infected cells, which corresponds roughly to 2x the number of double infected cells.

Code for this analysis can be found in *finddoubles.m*.

#### Single cell analysis

Code for analysis of single cells sequencing data can be found in *preprocessing_singlecells.sh*, *matlab_preprocessing_singlecells.m* and *analyse_singlecells.m.*

#### False positive rate

Code for the calculation of the false positive rate can be found in *analyse_bilateralinjections.m.* Dimensionality reduction and clustering.

Code for t-SNE dimensionality reduction and hierarchical clustering of cortico-cerulear neurons can be found in *doclusteringm.*

### Supplementary Note 5: Spike-in recovery

To assess the efficiency of barcode recovery in MAPseq, we added a known amount of spike-in RNA (Supplementary Fig. 1h) into every sample (Fig. 4a, Supplementary Fig. 3) and quantified the number of distinct spike in molecules in the sequencing results. The ratio of the number of recovered spike-in molecules to the number of input molecules then is the probability of detection of any given barcode molecule.

Detection efficiencies are relatively constant across areas and animals (Supplementary Fig. 7c,d) and average to P(detection)=0.024 for target areas. This implies that when we do not detect a barcode in an area, there are less than 123 barcode mRNA molecules present in that sample with a confidence >95%, as dictated by the negative binomial distribution.

Note here, that this measure of barcode detection probability is based on the efficiency of going from total RNA to sequencing results. It is blind to losses incurred during extraction of total RNA from tissue, such that the overall MAPseq detection efficiency is likely somewhat lower than what we estimate.

### Supplementary Figures

**Supplementary Figure 1;.**
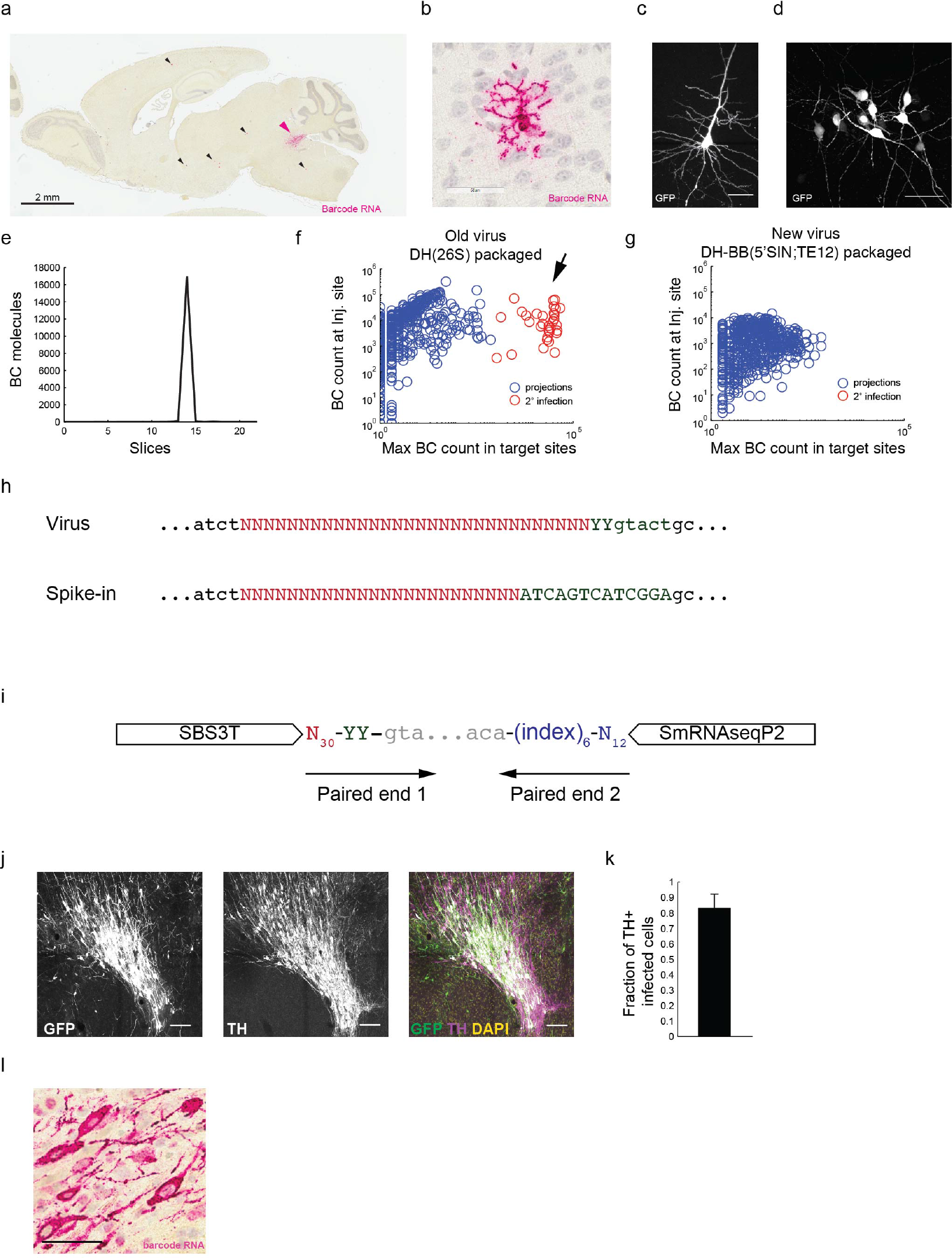
Related to Fig. 1 and Fig 2 Barcodes are delivered to LC neurons using recombinant Sindbis virus, (a-g) The replacement of the conventional packaging system, DH(26S)5’SIN, with a modified packaging system we developed, DH-BB(5’SIN;TE120RF), largely eliminates infection of cells distal to the injection site. After injection of conventionally packaged virus, (a,b) *in situ* hybridization for barcode mRNA labels cells far away from the injection site. Pink arrow=primary injection site; black arrows=secondary infection. (c,d) Similarly we can detect GFP positive cells or clusters of cells far away from the injection site after injection of conventionally produced virus. Scale bar=50μm. (e) MAPseq data produced using DH(26S)5’SIN packaged virus shows spurious barcodes with extremely high abundance in a single target site only (“spikes”), which arise from barcodes expressed in cortical somata secondarily infected by propagation of viral particles from the axons of infected LC neurons. (f) Expression levels of these high abundance barcodes are comparable to that of barcodes in the injection site. (g) Changing the packaging system to the new DH-BB(5’SIN;TE12ORF) produces a propagation incompetent Sindbis virus and eliminates these high abundance barcodes. All MAPseq results described in this manuscript made use of this new virus. (h) Differences in the sequence of viral barcodes and spike-in RNA allow easy discrimination of the two. (i) Structure of the final sequencing amplicon. (j,k,l) Stereotaxic injection of Sindbis virus reliably infects LC and fills cell bodies and axons with barcode mRNA. (j) Maximum z-projection of a representative Sindbis injection shows excellent overlap with the TH-stained LC, confirming successful stereotactic targeting of the nucleus. Scale bar=100μm. (k) Quantification of the fraction of infected cells that are also TH+ confirms reliable targeting of LC by streotactic injection. N=6. Mean +/- s.d. is shown. (l) RNA *in situ* of barcode mRNA showing good fills of cell bodies at the injection site. Scale bar=50μm.Number of infectedcells Sequence rank

**Supplementary Figure 2;.**
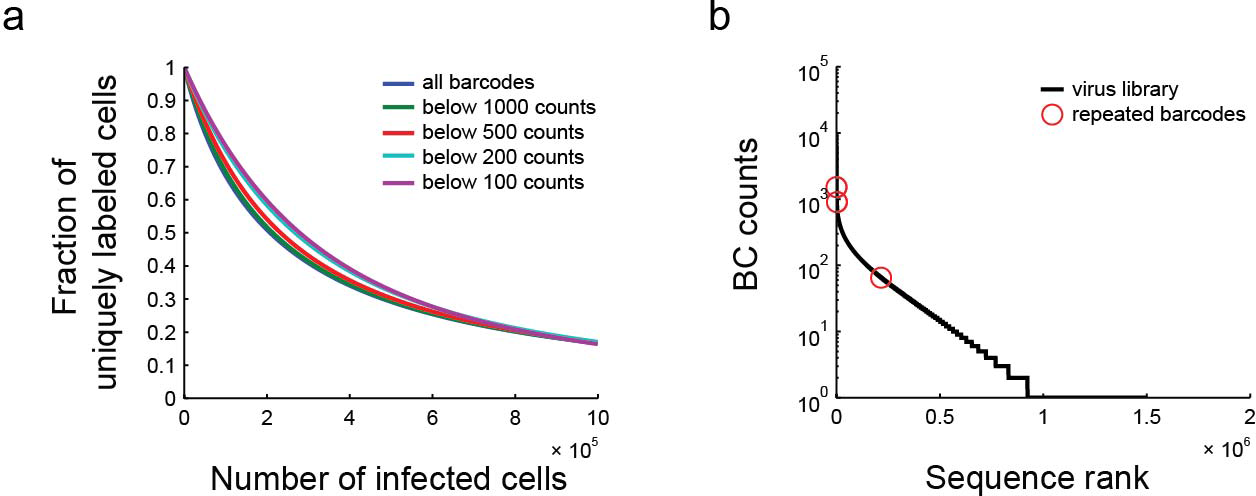
Related to Fig.3 The diversity of the MAPseq virus library is sufficient to uniquely label many cells, (a) The number of cells that can be uniquely labeled using our virus library does not change dramatically when we bioinformatically remove overrepresented barcodes from the library. The legend indicates which barcodes are still considered for labeling, (b) Position of the three barcodes that were traced in more than one of four animals. Two of the three are highly abundant in the virus library.

**Supplementary Figure 3;.**
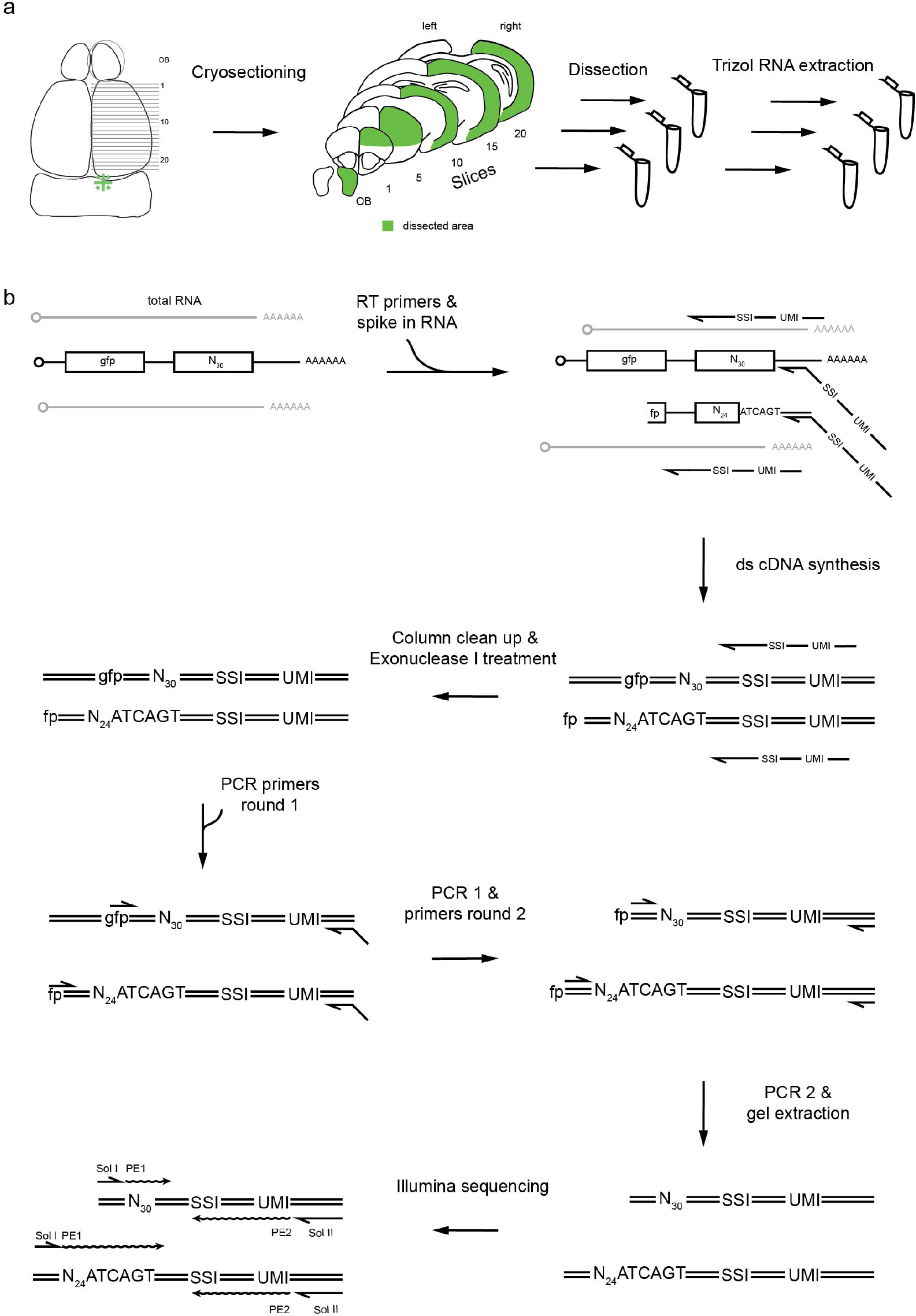
Related to Fig. 4 MAPseq workflow, (a) We cryosection a flash frozen brain and dissect out areas of interest. We then extract total RNA from every area individually, (b) To the total RNA from every area, we add a known amount of spike-in RNA and reverse transcription primers containing unique SSIs and UMIs. We produce double stranded cDNA, and digest leftover reverse transcription primers using Exonuclease I to avoid UMI containing primers to participate in subsequent PCR reactions. We then perform two rounds of nested PCR, bringing in the PE2 sequencing primer binding site and P7 sequence as5’ overhangs of the reverse primer. After gel extraction, the amplicons are ready for Illumina sequencing.

**Supplementary Figure 4;.**
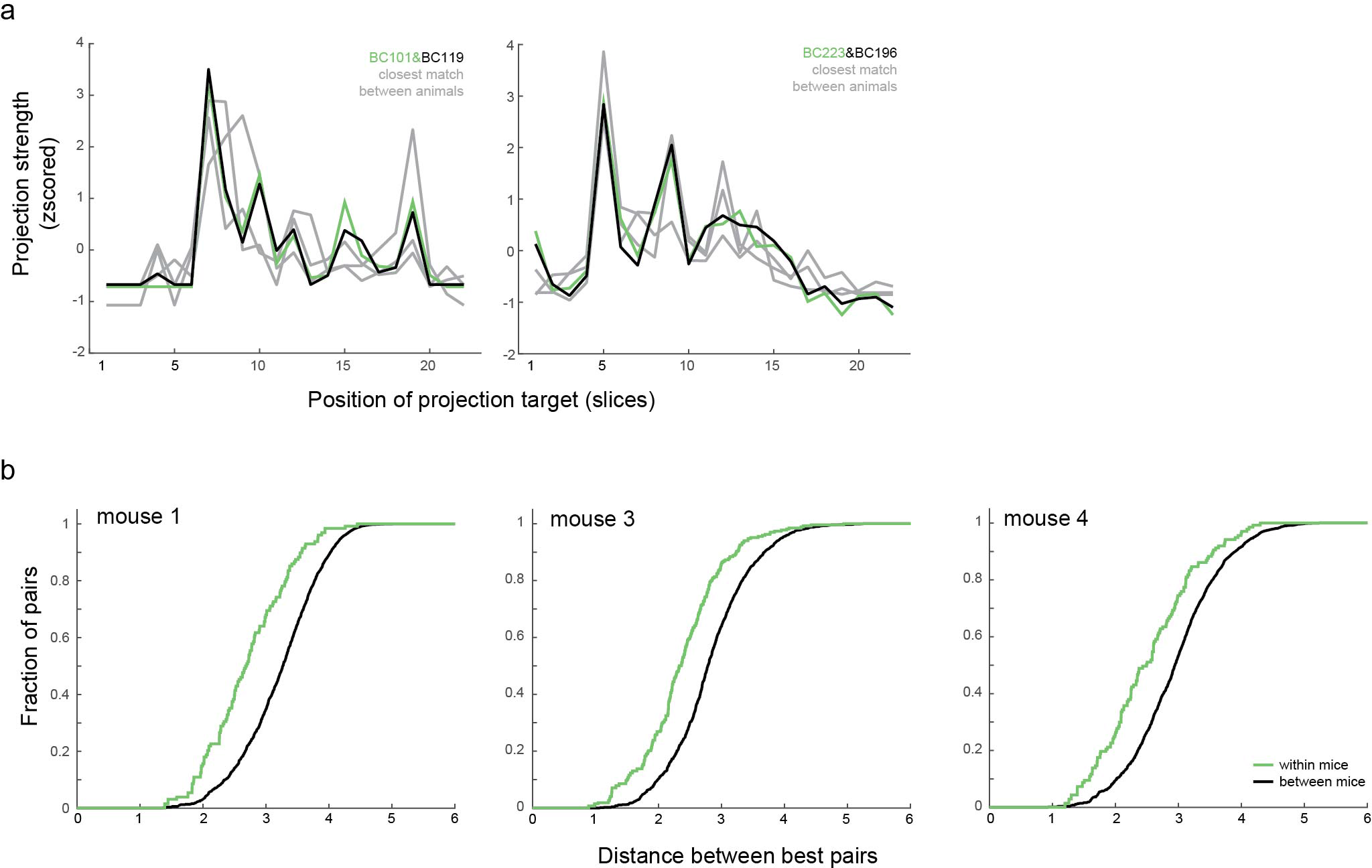
Related to Fig. 5 MAPseq provides a robust readout of single neuron projection patterns, (a) The same example pairs of barcodes profiles that are more similar than expected by chance as shown in Fig 5a. In grey we indicated best matches of the barcode profiles across animals from 5 independent samplings of the comparison animal, (b) Cumulative distribution function of distances of best barcode pairs within and across animals for animals 1, 3 and 4.

**Supplementary Figure 5;.**
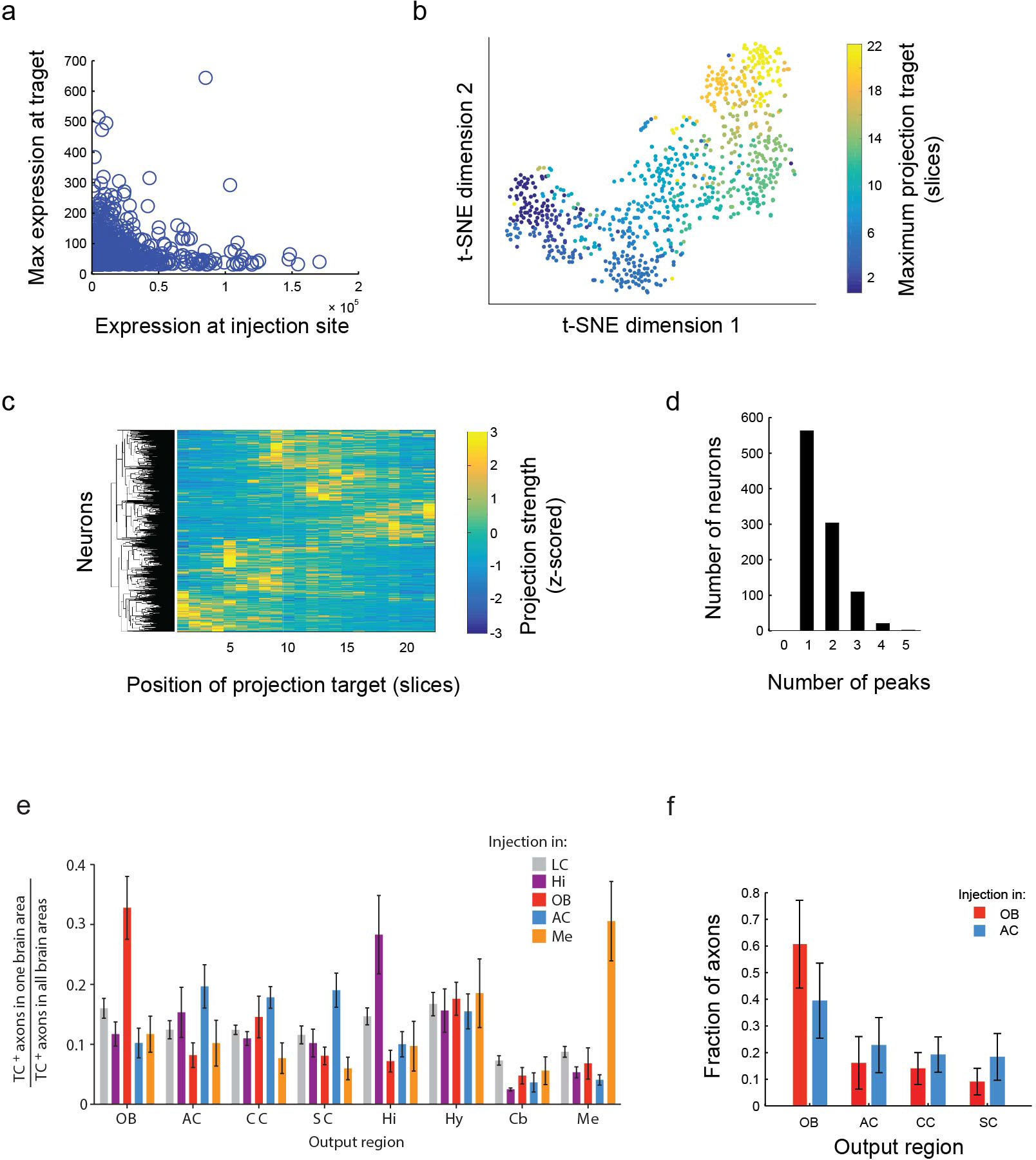
Related to Fig. 6 Aggregate projection of MAPseq traced neurons reproduces homogeneous bulk projection, but individual projection patterns are non-homogeneous. (a) There is no correlation between the expression level of a barcode at the injection site and its maximum projection strength to a target area, (b) t-SNE dimensionality reduction of cortically projection LC neurons reveals an orderly separation of neurons according to their maximum projection location, (c) Hierarchical clustering of z-scored projection profiles of cortical projection neurons however reveals no striking clustering, (d) Histogramof the number of detected peaks for all MAPseq traced neurons. For peak definitions see Supplementary Note 4. (e,f) Simulation of CAV-cre injection and axon tracing from MAPseq data reproduces the non-specific output pattern of LC neurons reported by Schwarz *et al.* (Schwarz et al., 2015). (e) Reproduction of Figure 4d of ref (Schwarz et al., 2015). Briefly, Schwarz *et al.* injected retrograde CAV-cre virus into a number of areas including olfactory bulb and auditory cortex, and ere dependent TVA-mCherry-AAV into LC.

They then counted the number of mCherry labeled LC axons in a number of output areas and normalized the number of axons across all output areas. They could thereby quantify the projection strength of groups of LC neurons defined by their projection to the injection site and found that most groups of LC neurons project equally to all output areas. (f) Results of our MAPseq data based simulation of the experiment preformed by Schwarz *et al,* plotted in the same way. Briefly, we simulated CAV-cre injections into olfactory bulb or auditory cortex by labeling barcodes that are present at more than 50 counts in either olfactory bulb or auditory cortex. We then summed up the normalized counts of the labeled barcodes in slices containing the output regions and normalized the resulting projection strength across all output regions, thus mimicking the counting of labeled axons in output regions. In contrast to the idiosyncratic single cell projection pattern reported by MAPseq, this simulation recapitulates the findings of Schwarz *et al.,* highlighting the importance of single neuron resolution in connectivity mapping. OB=olfactory bulb; AC=auditory cortex; CC=cingulate cortex; SC=somatosensory cortex.

**Supplementary Figure 6;.**
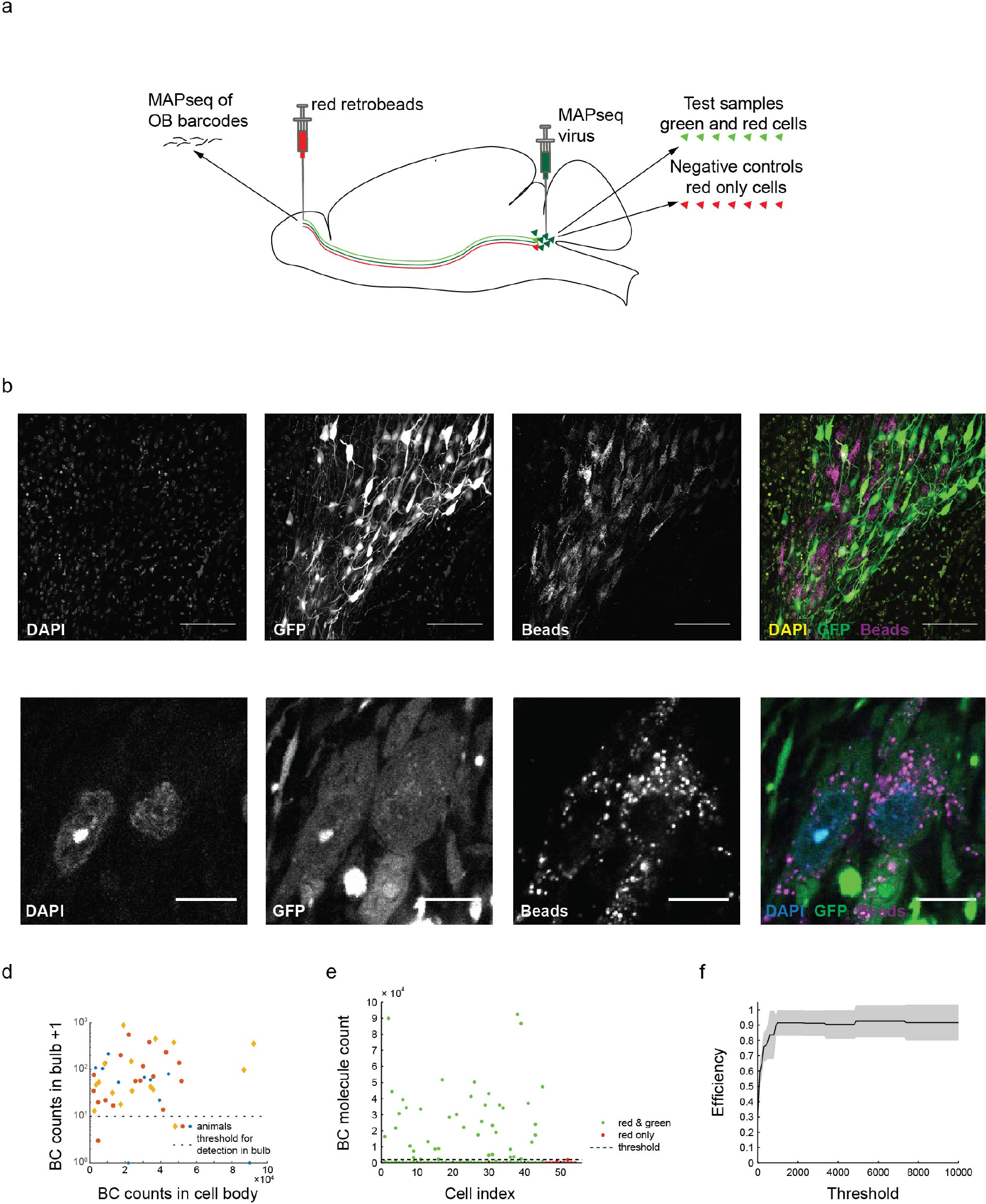
Related to Fig. 6 Sequencing of single LC cells reveals low MOI and high MAPseq efficiency, (a) Overview of the experimental design. Red Lumafluor retrobeads label bulb projecting cells in LC. Barcodes present in these cells should also be present in the bulb, (b) Overview image of LC, showing bead and GFP labeling of cells. Scale bar=lOOμm. (c) Detailed image of retrobeads andGFP labeled cells. Scale bar=10μm. (d) Scatter plot showing the relationship of barcode abundance in the olfactory bulb to barcode abundance in individual cells. The dashed line indicates the minimum barcode abundance in the bulb chosen as detection threshold. (e) Scatter plot of abundance of all barcodes found in the sequenced single cells for both bead and Sindbis labeled cells (n=45 from 3 animals, green) and negativecontrol cells (bead labeled only; n=9 from 3 animals; red). Dotted line indicates the height of the most abundant barcode from red only cells, the threshold chosen to distinguish real from artefactual barcodes. (f) MAPseq efficiency as a function of an increasingly stringent noise threshold. The MAPseq efficiency estimate is not very sensitive to changes in the threshold value. Shaded area indicates s.d. across animals.

**Supplementary Figure 7;.**
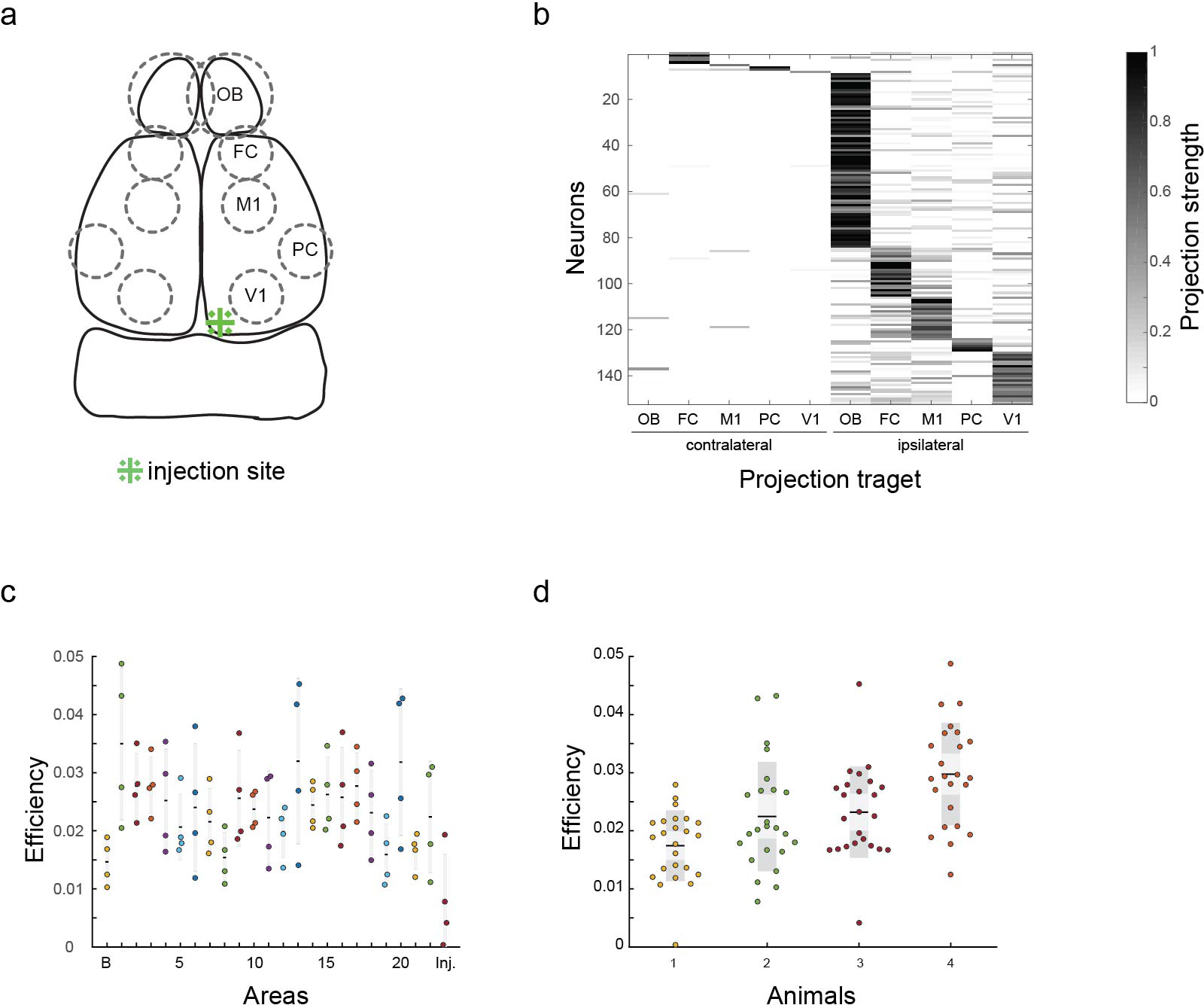
Related to Fig. 6 MAPseq can be performed on small target areas, (a) Schematic of dissected areas. FC=frontal cortex; Ml=primary motor cortex; PC=piriform cortex; VI=primary visual cortex, (b) A heatmap of all ~140 neurons traced across 3 independent animals using DH(26S)5’SIN packaged MAPseq virus. We removed all ectopically infected cells (see Supplementary Fig. 1f) that could have confused tracing results by a maximum abundance cutoff of 1000. Preferential targeting of different ipsilateral areas is clearly evident. (c,d) Efficiency of barcode recovery from total RNA samples in MAPseq is low and relatively constant across areas (c; n=4 animals) and animals (d) as measured by spike-in RNA recovery.

